# Divergent effects of absolute evidence magnitude on decision accuracy and confidence in perceptual judgements

**DOI:** 10.1101/2021.07.04.451079

**Authors:** Yiu Hong Ko, Daniel Feuerriegel, William Turner, Helen Overhoff, Eva Niessen, Jutta Stahl, Robert Hester, Gereon R. Fink, Peter H. Weiss, Stefan Bode

**Affiliations:** Melbourne School of Psychological Sciences, The University of Melbourne, Australia; Cognitive Neuroscience, Institute of Neuroscience and Medicine (INM-3), Research Centre Jülich, Germany; Department of Psychology, Faculty of Human Sciences, University of Cologne; Department of Neurology, University Hospital Cologne and Faculty of Medicine, University of Cologne

**Keywords:** decision-making, confidence, metacognition, change-of-mind, perception, absolute evidence

## Abstract

Whether people change their mind after making a perceptual judgement may depend on how confident they are in their decision. Recently, it was shown that, when making perceptual judgements about stimuli containing high levels of ‘absolute evidence’ (i.e., the overall magnitude of sensory evidence across choice options), people make less accurate decisions and are also slower to change their mind and correct their mistakes. Here we report two studies that investigated whether high levels of absolute evidence also lead to increased decision confidence. We used a luminance judgment task in which participants decided which of two dynamic, flickering stimuli was brighter. After making a decision, participants rated their confidence. We manipulated relative evidence (i.e., the mean luminance difference between the two stimuli) and absolute evidence (i.e., the summed luminance of the two stimuli). In the first experiment, we found that higher absolute evidence was associated with decreased decision accuracy but increased decision confidence. In the second experiment, we additionally manipulated the degree of luminance variability to assess whether the observed effects were due to differences in perceived evidence variability. We replicated the results of the first experiment but did not find substantial effects of luminance variability on confidence ratings. Our findings support the view that decisions and confidence judgments are based on partly dissociable sources of information, and suggest that decisions initially made with higher confidence may be more resistant to subsequent changes of mind.

## Introduction

The cognitive and neural processes underlying simple decisions have been studied extensively over the past decades, and performance in discrete choice tasks has been successfully accounted for using computational models (Gold & Shadlen, 2007; Smith & Ratcliff, 2004; Ratcliff et al., 2018). The most prominent class of models are evidence accumulation models, such as the Diffusion Decision model (DDM; Ratcliff, 1978). These models describe the decision process as a noisy accumulation of evidence towards alternative decision thresholds. For a discrete choice perceptual decision, such as deciding whether a cloud of dots are predominantly moving to the left or the right, these models propose that sensory evidence is sampled and integrated over time, and a decision is made when the accumulated evidence reaches a threshold in favour of a particular choice outcome.

When an incorrect decision is made, we can often rapidly detect that an error has occurred (Ullsperger et al., 2014). For example, in a typical Flanker task, when judging the identity of a central letter in the presence of distracting flankers, a detected decision error is reflected in brain activity following the incorrect motor response (Scheffers & Coles, 2000). Beyond simply detecting an error, we can also rapidly change our minds and correct erroneous decisions (Resulaj et al., 2009; van den Berg et al., 2016). This capacity for fast changes of mind has been linked to metacognitive processes – specifically, those which allow us to derive a subjective sense of confidence in our decisions (van den Berg et al., 2016; Fleming & Daw, 2017). Consistent with the notion of a link between the processes underlying confidence and changes of mind, decisions initially made with high confidence are less likely to be overruled than those made with a lower degree of confidence (van den Berg et al., 2016). In light of this, it has been proposed that decision confidence may influence the processes that determine how quickly and how often we change our minds (Turner, Feuerriegel, et al., 2021; van den Berg et al., 2016). Beyond these initial proposals, however, this idea is yet to be systematically tested.

### The effect of absolute evidence on changes of mind

A recent study investigated how variations in absolute evidence magnitude affect the speed and accuracy of decisions and subsequent changes of mind (Turner, Feuerriegel et al., 2021). In this study, participants judged which of two flickering squares was, on average, brighter by integrating information over a short period of time. The overall (i.e., summed) luminance across the two stimuli was manipulated to investigate the effect of absolute evidence, while relative evidence (i.e., their luminance difference) was held constant. According to certain classes of evidence accumulation models (i.e., ‘purely relative’ models such as the DDM), the *differences* in evidence for each choice option are accumulated in the decision process (Ratcliff & Rouder, 1998). This is empirically supported by the standard finding that increasing relative evidence leads to higher accuracy and faster response times (RTs; Ratcliff et al., 2018; Teodorescu et al., 2016). However, purely relative models do not predict an effect of absolute evidence on decision accuracy or RT if relative evidence is held constant (although performance differences at different absolute evidence levels could still emerge due to Weber-scaling, i.e., diminished increase in perceived brightness as luminance increases; Geisler, 1989). Nevertheless, Turner, Feuerriegel and colleagues (2021) showed that, even after considering the effects of Weber-scaling, decisions are sensitive to variations in absolute evidence magnitude – a finding which purely relative models cannot account for. In particular, consistent with other studies, it was shown that increasing absolute evidence leads to faster but less accurate decisions (Ratcliff et al., 2018; Teodorescu et al., 2016; Turner, Feuerriegel, et al., 2021).

It should be noted that, to be consistent with previous studies (Peters et al., 2017; Teodorescu et al., 2016; Turner et al., 2021), we use the term ‘evidence’ to refer to sensory evidence. That is, sensory information about each of the choice options. Therefore, in concrete terms, we define the ‘evidence’ for each choice option as their respective luminance values. Importantly, under this definition, the term ‘evidence’ should not be interpreted as necessarily referring to ‘choice evidence’. That is, information which can be used to inform a choice. This is because the sensory evidence associated with a single choice option (i.e., the luminance of one of the squares), or indeed the overall level of absolute evidence, is by itself not informative for decision-making (at least for comparative judgements).

Turner, Feuerriegel, and colleagues (2021) asked the additional question of how absolute evidence magnitude affects the speed and likelihood of change-of-mind decisions. In their study, the stimuli were first presented for an initial luminance judgment and then remained on the screen for a further 1 s, allowing participants to submit a second, change-of-mind response within this time window. They reported that higher levels of absolute evidence led to slower change-of-mind RTs relative to the time of the decision. Importantly, these RT effects also remained when effects of Weber-scaling were accounted for in a follow-up experiment. This finding suggests that participants may have required a larger amount of conflicting, post-decisional sensory evidence to overrule their decisions in conditions of high absolute evidence. As decision RTs were consistently faster in higher absolute evidence conditions, and faster RTs are associated with higher levels of decision confidence (Kiani et al., 2014), participants may have been more confident in their decisions, despite being objectively less accurate. If this were the case, this might have led participants to wait longer and accumulate more evidence before deciding to overrule their decision. The current study aimed to examine whether confidence would increase with increased absolute evidence. This would point to a moderation effect of confidence that could ultimately drive changes of mind. We further tested more directly whether any effects of confidence would potentially translate into changes of mind by converting our confidence measure into a change-of-mind measure, following previous approaches (Charles & Yeung, 2019; Fleming et al., 2018).

### The decision-congruent evidence hypothesis

The idea that confidence may have increased with higher absolute evidence magnitude is consistent with the *decision-congruent evidence hypothesis*. This hypothesis suggests that the extent of sensory evidence in favour of the selected option primarily informs confidence judgments (Koizumi et al., 2015; Odegaard et al., 2018; Peters et al., 2017; Samaha & Denison, 2020; Zylberberg et al., 2012). These accounts suggest that, while decisions are determined by the difference in evidence between choice options (i.e., relative evidence), confidence might be a product of a winner-takes-all process. The more evidence for the winning option, the higher the confidence in the decision, regardless of the amount of evidence for the alternative, non-chosen option (Peters et al., 2017; Zylberberg et al., 2012). Similar results have been shown experimentally via manipulations of ‘positive’ evidence (i.e., evidence supporting the correct response) and ‘negative’ evidence (i.e., evidence supporting the alternative response) in random dot motion and grating orientation judgment tasks (Koizumi et al., 2015; Odegaard et al., 2018; Samaha et al., 2016; Samaha & Denison, 2020). Specifically, when the ratio between positive and negative evidence was held constant, increased positive and negative evidence together led to increased confidence without impacting accuracy. This effect was termed the *Positive Evidence Bias* (PEB; Maniscalco et al., 2021; Samaha & Denison, 2020). Taken in relation to the findings of Turner, Feuerriegel, and colleagues (2021), this would mean that stronger absolute evidence (and the corresponding increase in decision-congruent evidence) may have increased participants’ subjective confidence in their decisions and, in turn, made them less prone to changing their mind. As Turner, Feuerriegel and colleagues (2021) did not investigate confidence, the current study was designed to directly test whether stronger absolute evidence in the same task as used by Turner and colleagues does indeed lead to increased decision confidence.

### The current study

We employed a luminance discrimination task using flickering stimuli, as in Turner, Feuerriegel, and colleagues (2021), and manipulated both absolute and relative evidence across three levels (low, medium, and high). Stimuli were presented for a maximum of 1.5 s and disappeared when the keypress response reported the perceptual decision. Participants subsequently reported their degree of confidence in their decision on a 7-point scale ranging from “surely incorrect” to “surely correct”.

In Experiment 1, we first aimed to replicate previous findings that (i) increasing relative evidence leads to increased decision accuracy and faster RTs, and (ii) increasing absolute evidence leads to both lower accuracy and faster RTs (Ratcliff et al., 2018; Teodorescu et al., 2016; Turner, Feuerriegel, et al., 2021). Furthermore, we predicted that confidence in trials with correct responses would increase with stronger relative evidence, while confidence in error trials would decrease with stronger relative evidence, as shown in previous studies (Sanders et al., 2016). Critically, we also predicted that stronger absolute evidence would be associated with increased confidence for both correct and error trials, despite decreased decision accuracy. This is because higher absolute evidence implies stronger decision-congruent evidence for both correct and error trials. Such findings in a highly similar task to Turner, Feuerriegel et al. (2021) would suggest that slower change-of-mind responses co-occur with a higher degree of confidence in one’s decision.

It should be taken into account that the high absolute evidence stimuli (i.e., brighter pairs of squares) would likely have been perceived as being less variable over time. This is because physical luminance and perceived brightness are related via a nonlinear compressive function (Geisler, 1989). In other words, under conditions of high luminance, changes in perceived brightness with an equivalent increase in luminance are diminished. Accordingly, when there is the same amount of luminance variability, the variability in brightness over time will be perceived as more pronounced in the dimmer stimulus condition (i.e., dimmer squares will appear to flicker more than brighter squares).

Perceived stimulus variability is theorised to inform confidence judgements (Yeung & Summerfield, 2012). Consistent with this theory, some previous studies have shown that higher stimulus variability can lead to lower confidence ratings (e.g., Spence et al., 2016; Navajas et al., 2017; Desender et al., 2018). However, the opposite has also been found (e.g., Zylberberg et al., 2014, 2016), where the observed effects appear to vary by the task and type of variability manipulation. It is therefore possible that reductions in perceived brightness variability for higher absolute evidence conditions in our task might have led to higher confidence ratings, which might explain the observed effects of absolute evidence.

In Experiment 2, we examined whether decreases in stimulus variability could lead to increased confidence in our experimental design, by directly manipulating luminance variability (i.e., the distribution of luminance values between frames around the same mean), in addition to relative and absolute evidence. This experiment therefore served two purposes: a) to replicate the general effects of Experiment 1, and b) to directly (i.e., experimentally) test the effect of stimulus variability on confidence within our specific luminance task. While finding that reduced stimulus variability leads to substantially higher confidence ratings would not prove that this is indeed the explanation for why increased absolute evidence impacts confidence, it would nevertheless demonstrate that it is possible for stimulus variability to affect confidence in our study design. We could then speculate that this might also be a potential alternative explanation for the effects of absolute evidence on confidence in our experiments. However, if directly manipulating stimulus variability does not affect confidence in our study, a relevant contribution of stimulus variability on the current findings can be essentially ruled out. Moreover, we could include stimulus variability in our models to test whether effects of absolute evidence are reproducible at different levels of stimulus variability.

## Experiment 1

### Methods

#### Participants

Thirty-seven university student volunteers with normal or corrected-to-normal vision were recruited. Six participants were excluded: Three failed to report confidence in more than 20% of all trials, two showed lower than 55% accuracy, and one reported the same confidence level in more than 90% of trials where confidence was reported. The final sample comprised 31 participants (mean age = 26 years, SD = 5, range 19 – 38 years, 17 females). This experiment was approved by the University of Melbourne ethics committee (ID: 1954641.2).

#### Experimental Procedures

Before the experiment, participants gave written consent and were given task instructions. Participants were then seated in a dark testing booth 70 cm from a computer monitor. At the beginning of the experiment, participants underwent task training while the experimenter stayed in the testing booth. This procedure ensured that participants understood the task instructions correctly. Participants then completed the main experiment. After completion of the task, participants were reimbursed 20 AUD and were debriefed by the experimenter.

#### Task and Stimuli

We used a luminance judgment task to examine the effects of relative and absolute evidence on perceptual decisions, RTs, and decision confidence. Participants had to decide which of two flickering squares, presented on the left and right of a central red fixation dot, was brighter (Figure 1A). There were three levels (low, medium, high) for both relative evidence and absolute evidence, resulting in a 3 × 3 factorial design (Figure 1B).

**Figure 1.**
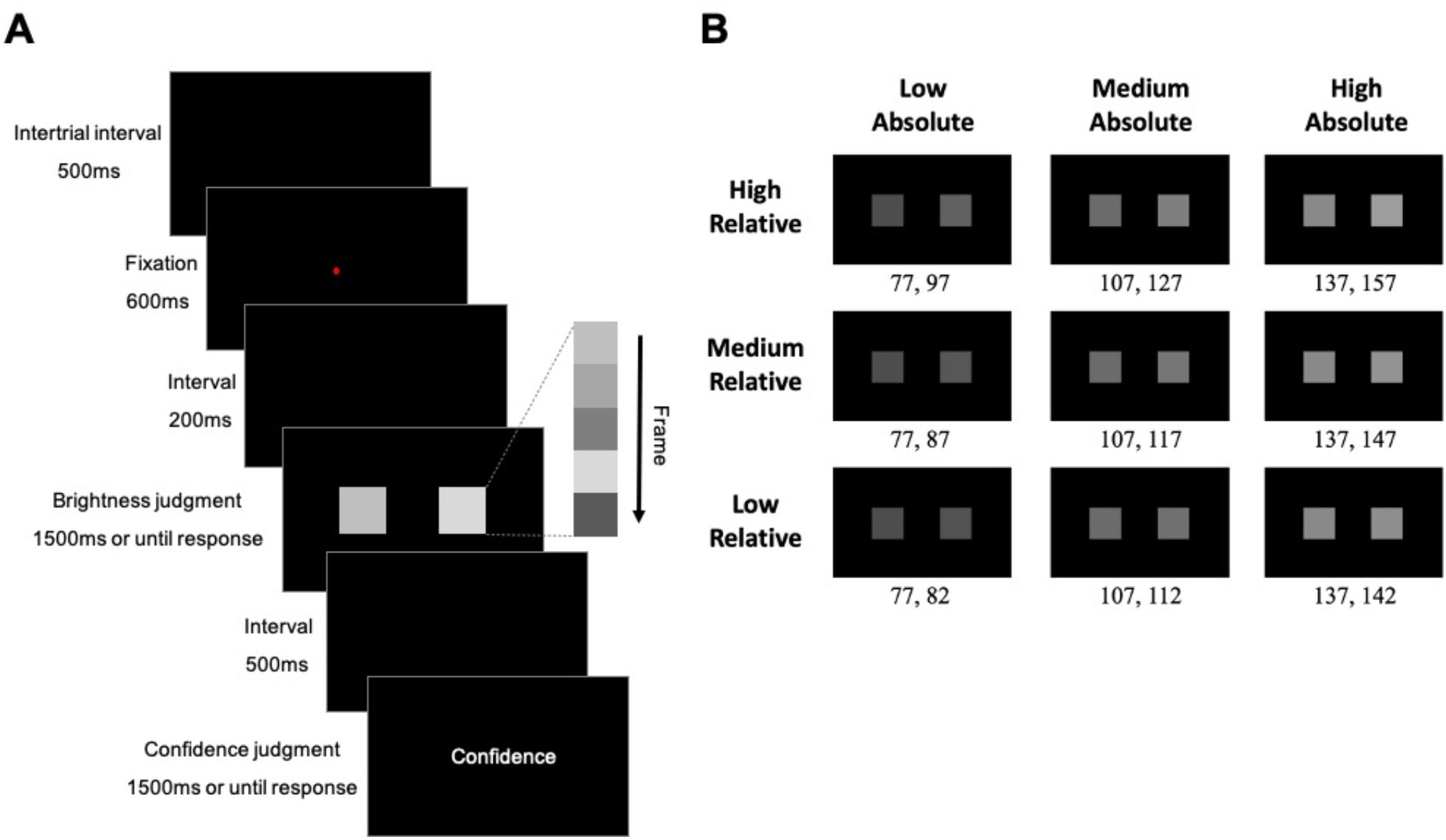
Task paradigm and stimuli. (A) Paradigm. In each trial, two flickering square stimuli of different average luminance were presented. Each square changed in luminance with each frame. Participants were required to select the stimulus that appeared brighter on average and subsequently reported their decision confidence using a 7-point scale while the word “confidence” was presented on the screen. (B) Illustration of average luminance values for stimuli for all experimental conditions of Experiment 1. Luminance values were randomly sampled from normal distributions truncated one standard deviation around pre-defined means. The standard deviation of all distributions was 25.5.

The two square stimuli changed in luminance with each frame refresh (i.e., every 13.3 ms at 75 Hz). For each refresh, the luminance value for each square was determined by randomly drawing values sampled from two truncated normal distributions around the pre-determined means for each square, respectively. Mean luminance values for the two distributions were specified by pairs of RGB values, such that one distribution had a higher mean than the other (therefore, one stimulus appeared on average brighter than the other; Figure 1B). Following Ratcliff et al. (2018), both distributions had a standard deviation of 25.5 and were truncated at ±1 SD from their means. The mean luminance values mapped onto relative evidence strength, defined as the difference in distribution means for the two stimuli, and absolute evidence strength, defined as the sum of the distribution means for both stimuli. The size of both stimuli was 70 x 70 pixels, and they were positioned at equal distance from the centre of the screen, separated from each other by 180 pixels. The positions of the stimuli were counterbalanced such that in half of the trials, the left stimulus was brighter, and in the other half of the trials the right stimulus was brighter. The order of the trials was randomised. Participants were instructed to respond as quickly as possible. After submitting the choice response, participants were required to indicate their confidence using a 7-point rating scale ranging from “surely incorrect” (1) to “surely correct” (7), with a midpoint rating (4) indicating they were unsure whether they were correct or incorrect (i.e., they felt they were guessing). They were again instructed to respond as fast as possible.

Participants completed a training phase before starting the experimental phase. They first practised the experimental task (see below) for 32 trials, without making confidence ratings. Instead, they received performance feedback after each trial to familiarise themselves with the judgement task. Subsequently, they practised the entire task, including confidence judgements for another 32 trials in which (as in the main experiment) no performance feedback was given. During training, a confidence rating scale was presented on the screen for 500 ms. Participants were instructed that during the main task, this visual presentation of the scale would be removed, and only the word “confidence” would prompt the rating.

##### Experiment

After the two training blocks, participants started the main experiment. Each trial started with an intertrial interval lasting 500 ms. A red fixation dot followed this interval in the middle of the screen for 600 ms, and then a blank screen was presented for 200 ms. After that, the flickering squares were presented, and participants were required to make the brightness decision by pressing either the left or right key on a response pad using left and right index fingers, corresponding to which stimulus they perceived as brighter. The stimuli were presented for a maximum of 1,500 ms and disappeared immediately after a response was submitted. Subsequently, after an interval of 500 ms with a blank screen, participants were asked to rate their confidence without a visual presentation of a rating scale but prompted by the word “confidence”. The scale had the same properties as during training, and participants were required to press one of the seven keys on the response pad to indicate their confidence level. No confidence rating was required if the brightness judgment was “too slow” (>1,500 ms RT) or “too quick” (<250 ms RT). In this case, only the respective timing feedback was presented for 1,500 ms, and then the next trial began.

The experiment comprised 1,008 experimental trials equally allocated across 14 blocks. Each block was followed by a self-terminated rest period. An equal number of trials from all conditions were randomly interleaved within each block.

#### Apparatus

Stimuli were presented on a Sony Trinitron Multiscan G420 CRT Monitor (resolution 1280 x 1024 pixels; frame rate 75 Hz) that was gamma-corrected with a ColorCAL MKII Colorimeter (Cambridge Research Systems), such that the physical luminance of the stimuli was linearly related to the RGB values. The task was programmed in MATLAB R2018b (The Mathworks) using Psychtoolbox-3 (Brainard, 1997; Kleiner et al., 2007). Participants responded using a seven-button Cedrus response pad (RB-740, Cedrus Corporation).

#### Data Analysis

We used generalised linear mixed-effects models (GLMMs) to examine the effects of relative and absolute evidence on accuracy, RT, and confidence ratings. For RT and confidence ratings, we ran two separate sets of analyses: one included only correct trials and the second set included only error trials. This was done to control for the effect of accuracy, given that error trials have different RT distributions than correct trials, and confidence patterns for correct and error trials could also potentially differ across relative and absolute evidence conditions (Gajdos, Fleming, Saez Garcia, Weindel, & Davranche, 2019; Turner, Angdias, et al., 2021; Urai, Braun, & Donner, 2017). Additionally, by converting confidence ratings into a change-of-mind measure, we also analysed how change-of-mind frequency was affected by absolute and relative evidence strength. This was done by transforming confidence ratings into a binary variable (confidence lower than 4 as 1[change-of-mind trials], and confidence higher or equal to 4 as 0 [trials without changes of mind]), as in Charles and Yeung (2019) and Fleming et al. (2018). This approach resulted in seven separate and independent analyses with different dependent variables: accuracy, RT (correct), RT (error), confidence (correct), confidence (error), changes of mind (correct), and changes of mind (error).

For each model, the model structure included fixed effects of relative evidence, absolute evidence, and the interaction between relative and absolute evidence, and a random intercept by participant. We also attempted to fit models with random slopes for each effect of interest. However, we found that not all models with random slopes converged across the different analyses of accuracy, RT and confidence. To be consistent across these different analyses we therefore used models without random slopes.

As in Turner, Feuerriegel, and colleagues (2021), for different dependent variables different distributions were assumed, and different link functions were used: Binomial distributions with a logit function were used to model accuracy and changes of mind, gamma distributions with an identity function were used to model RTs, and normal distributions with an identity function were used to model confidence. All analyses were conducted in R (version 4.0.1). GLMMs were fitted using the lme4 package (version 1.1; Bates et al., 2015), and statistical significance of each effect was determined by likelihood ratio tests conducted using the afex package (version 0.28; Singmann et al., 2017). For brevity, only significant effects are reported in the results section. Complete statistical results including likelihood ratio test results for all effects and regression coefficients of the full models are reported in Supplementary Material. Code and data used for the analyses in this paper are available at https://osf.io/r8vfx/.

As shown by the results below, increases in absolute evidence led to increased confidence and faster RTs. As faster responding has been shown to contribute to higher confidence (Kiani et al., 2014), we further asked whether the effect of absolute evidence on confidence could simply be explained by faster RTs in conditions with higher absolute evidence. To answer this question, we ran post-hoc analyses in which confidence was predicted by RT and the same predictors as in the main analyses except absolute evidence, and then included absolute evidence in the model to examine whether absolute evidence could predict confidence above the effect of RT. Also similar to the main analyses, each variable was entered into the model in a forward stepwise approach, and the statistical significance of each predictor was determined by likelihood ratio tests comparing the models before and after the predictor was included.

### Results

#### Accuracy and Response Times

First, we aimed to replicate previous findings that accuracy increases with stronger relative evidence but decreases with stronger absolute evidence (Ratcliff et al., 2018; Teodorescu et al., 2016; Turner, Feuerriegel, et al., 2021). As expected, there was a positive effect of relative evidence (*χ²*[2] = 1433.01, *p* <.001), a negative effect of absolute evidence (*χ²*[2] = 485.87, *p* <.001), and an interaction (*χ²*[4] = 91.71, *p* <.001; the interaction was observed because the log odds of being correct was reduced by absolute evidence more strongly when relative evidence was high; see Supplementary Figure S1. However, this pattern was not observable in terms of proportion correct). Figure 2A shows that the average proportion of correct decisions increased with stronger relative evidence but decreased with stronger absolute evidence. Full statistical results are presented in Supplementary Tables S1 and S2.

**Figure 2.**
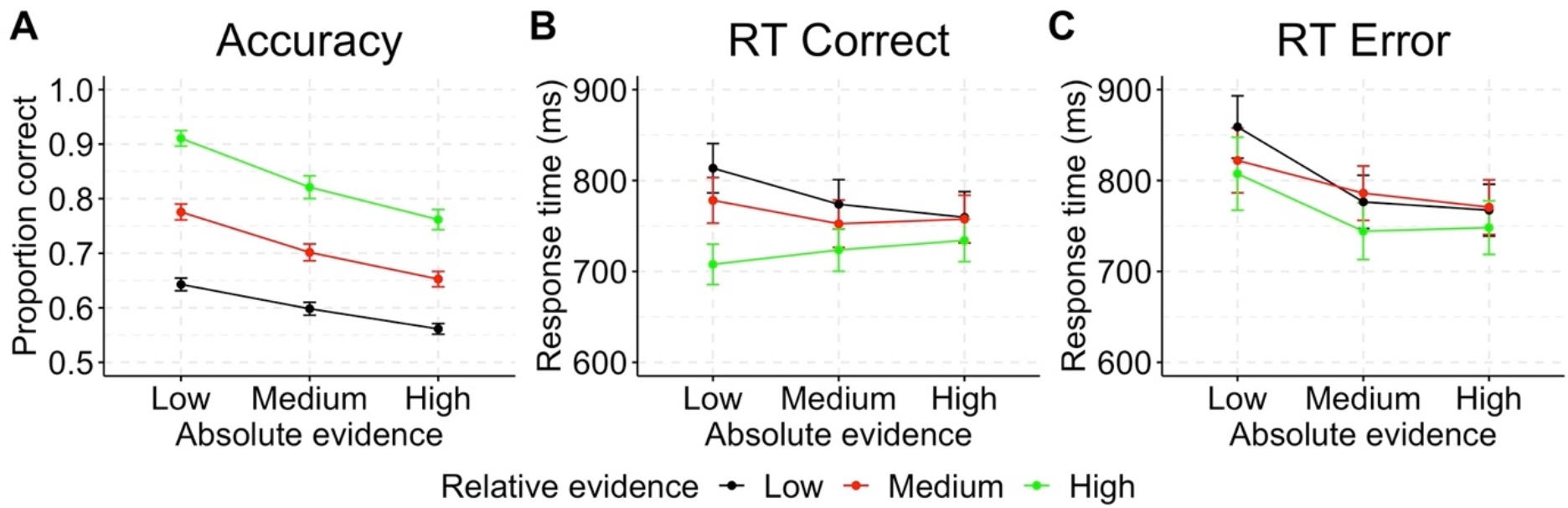
Experiment 1 accuracy and response time (RT). (A) Decision accuracy (average proportion correct) in each condition. (B) Mean RTs for correct trials. (C) Mean RTs for error trials. Error bars represent standard errors of the mean (SEM).

RTs were expected to be faster in conditions of stronger relative evidence and higher absolute evidence (Ratcliff et al., 2018; Teodorescu et al., 2016; Turner, Feuerriegel, et al., 2021). Consistently, for correct trials, there was an effect of relative evidence (*χ²*[2] = 314.71, *p* <.001), an effect of absolute evidence (*χ²*[2] = 35.85, *p* <.001), and an interaction (*χ²*[4] = 106.25, *p* <.001; Figure 2B). RTs were faster in conditions with stronger relative and stronger absolute evidence, and the effects of relative evidence appeared to diminish in conditions of higher absolute evidence. When the analysis was repeated for error trials only, similar effects were found (relative evidence: *χ²*[2] = 18.54, *p* <.001; absolute evidence: *χ²*[2] = 49.96, *p* <.001; interaction: *χ²*[4] = 20.20, *p* <.001; Figure 2C). Figure 2B and C show that RTs generally became faster with stronger relative evidence. Stronger absolute evidence also led to faster RTs for low and medium relative evidence, both for correct as well as error trials. The exception was that, for the high relative evidence condition, RTs were slower with low absolute evidence in error trials, but faster in correct trials. This result pattern was also reported in a previous study and appears to be a feature of this task (Ratcliff et al., 2018). Supplementary Figure S2 further shows that RTs were faster in conditions of higher absolute evidence across all RT quantiles, as also reported by Turner, Feuerriegel, et al. (2021). Full statistical results are presented in Supplementary Tables S3 – S6.

Taken together, these results show that increases in relative evidence are associated with faster and more accurate decisions. Moreover, these results replicate recent reports that increases in absolute evidence were associated with faster but less accurate decisions. The following section investigates the effect of absolute evidence magnitude on participants’ confidence ratings directly.

#### Confidence

We predicted that confidence would increase with both stronger relative and absolute evidence for correct trials. Consistent with our prediction, there was an effect of relative evidence (*χ²*[2] = 879.07, *p* <.001), an effect of absolute evidence (*χ²*[2] = 293.89, *p* <.001), and an interaction (*χ²*[4] = 121.55, *p* <.001). Figure 3A shows that mean confidence ratings increased with both relative and absolute evidence, although the effect of absolute evidence diminished as relative evidence increased. For the analysis of error trials, there was only an effect of relative evidence (*χ²*[2] = 99.99, *p* <.001) and an effect of absolute evidence (*χ²*[2] = 392.15, *p* <.001); Figure 3B). As expected, the direction of the relative evidence effect was reversed, with highest confidence ratings seen in lower as compared to higher relative evidence conditions. Full statistical results are presented in Supplementary Tables S7 – S10. Interestingly, even on error trials, absolute evidence magnitude was positively associated with confidence.

**Figure 3.**
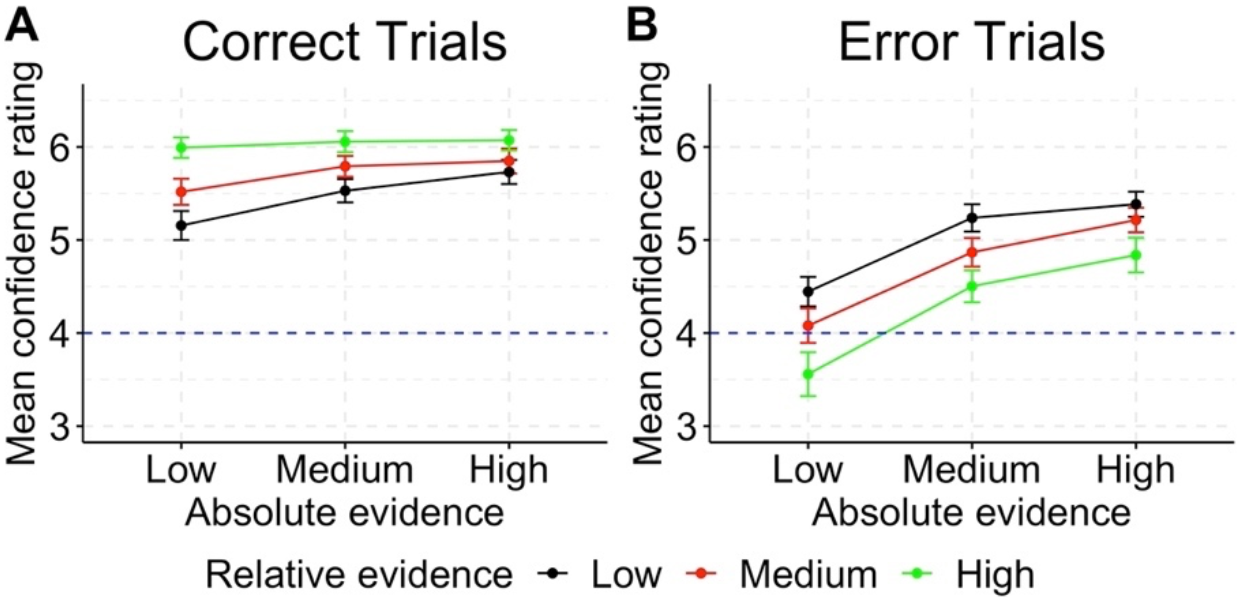
Mean confidence ratings in each condition in Experiment 1. (A) Correct trials. (B) Error trials. Confidence ratings were measured on a scale ranging from 1 (“surely incorrect”) to 7 (surely correct). The dotted line indicates the mid-point of the scale. Error bars represent SEM.

#### Change of Mind

When confidence was transformed into a binary variable that indicates changes of mind (confidence lower than 4 indicates a change of mind), in correct trials we observed a negative effect of relative evidence (*χ²*[2] = 259.51, *p* <.001), a negative effect of absolute evidence (*χ²*[2] = 38.67, *p* <.001), and an interaction between relative and absolute evidence (*χ²*[4] = 22.35, *p* <.001). Figure 4A showed that changes of mind were less likely with both stronger relative and absolute evidence, although the effect of absolute evidence diminished for stronger relative evidence. For error trials, a similar negative effect of absolute evidence was observed (*χ²*[2] = 176.64, *p* <.001) while relative evidence showed a positive effect (*χ²*[2] = 96.92, *p* <.001). There was also an interaction between relative and absolute evidence (*χ²*[4] = 14.09, *p* = .007). Figure 4B showed that changes of mind were less likely with stronger absolute evidence and weaker relative evidence, and the effect of absolute evidence was diminished when relative evidence was weaker. These patterns of results are opposite to the patterns of confidence, consistent with the negative relationship between confidence and changes of mind. Full statistical results are presented in Supplementary Tables S11 – S14.

**Figure 4.**
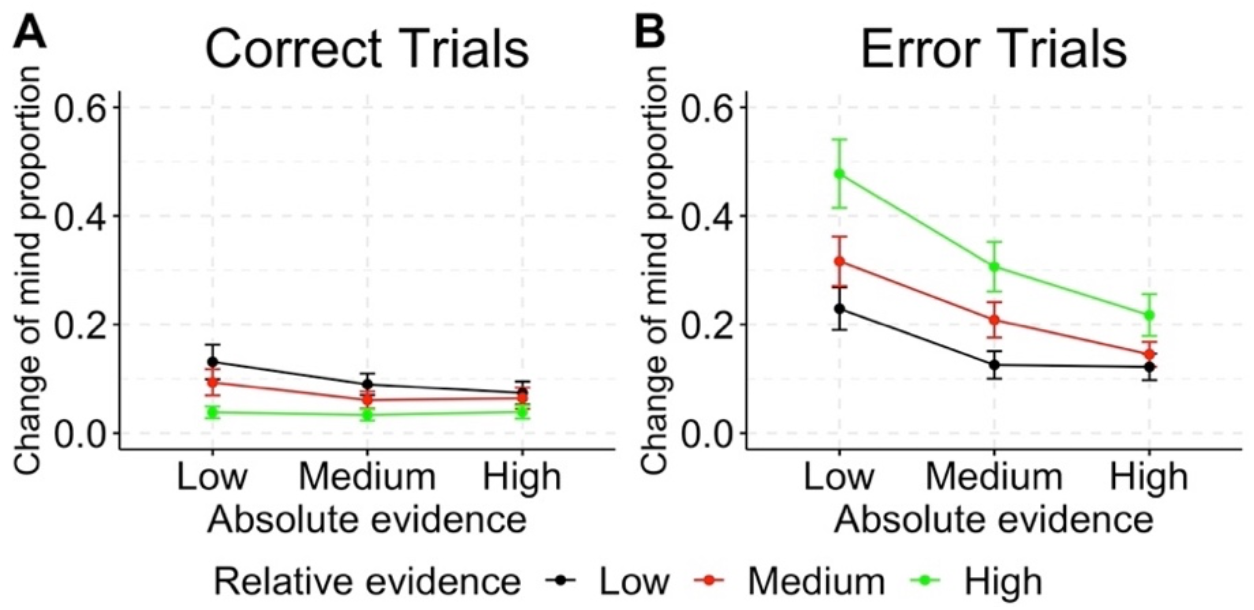
Experiment 1 proportions of change-of-mind trials in each condition. (A) Correct trials. (B) Error trials. Change-of-mind trials were defined as trials with confidence ratings lower than 4, indicating that the participant believed their brightness judgement was incorrect.

#### The Effect of Absolute Evidence on Confidence in Addition to RT

Lastly, to examine whether the effect of absolute evidence on confidence was simply due to faster RTs in higher absolute evidence conditions, we fitted models in which confidence was predicted by RT and relative evidence, and then compared model fits with a model that included the predictor of absolute evidence. When controlling for effects of RT in this way, confidence in correct trials was still predicted by relative evidence (*χ²*[2] = 667.60, *p* <.001) and RT (*χ²*[1] = 1207.10, *p* <.001), and additionally by absolute evidence (*χ²*[2] = 270.15, *p* <.001) as well as the interaction between relative and absolute evidence (*χ²*[4] = 80.22, *p* <.001). Similarly, confidence in error trials was also predicted by relative evidence (*χ²*[2] = 125.88, *p* <.001) and RT (*χ²*[1] = 405.39, *p* <.001), and additionally by absolute evidence (*χ²*[2] = 335.72, *p* <.001) and its interaction with relative evidence (*χ²*[4] = 10.37, *p* = .035). Full statistical results are presented in Supplementary Tables S15 – S18. To further visualize how confidence was related to RT and absolute evidence, we binned the data into six RT bins using RT quantiles of each participant (10%, 30%, 50%, 70%, 90%; separately for correct and error trials), and plotted mean confidence in each bin by absolute evidence in Figures 5A and B. These figures show that increasing absolute evidence generally led to higher confidence across correct and error trials across RT bins.

**Figure 5.**
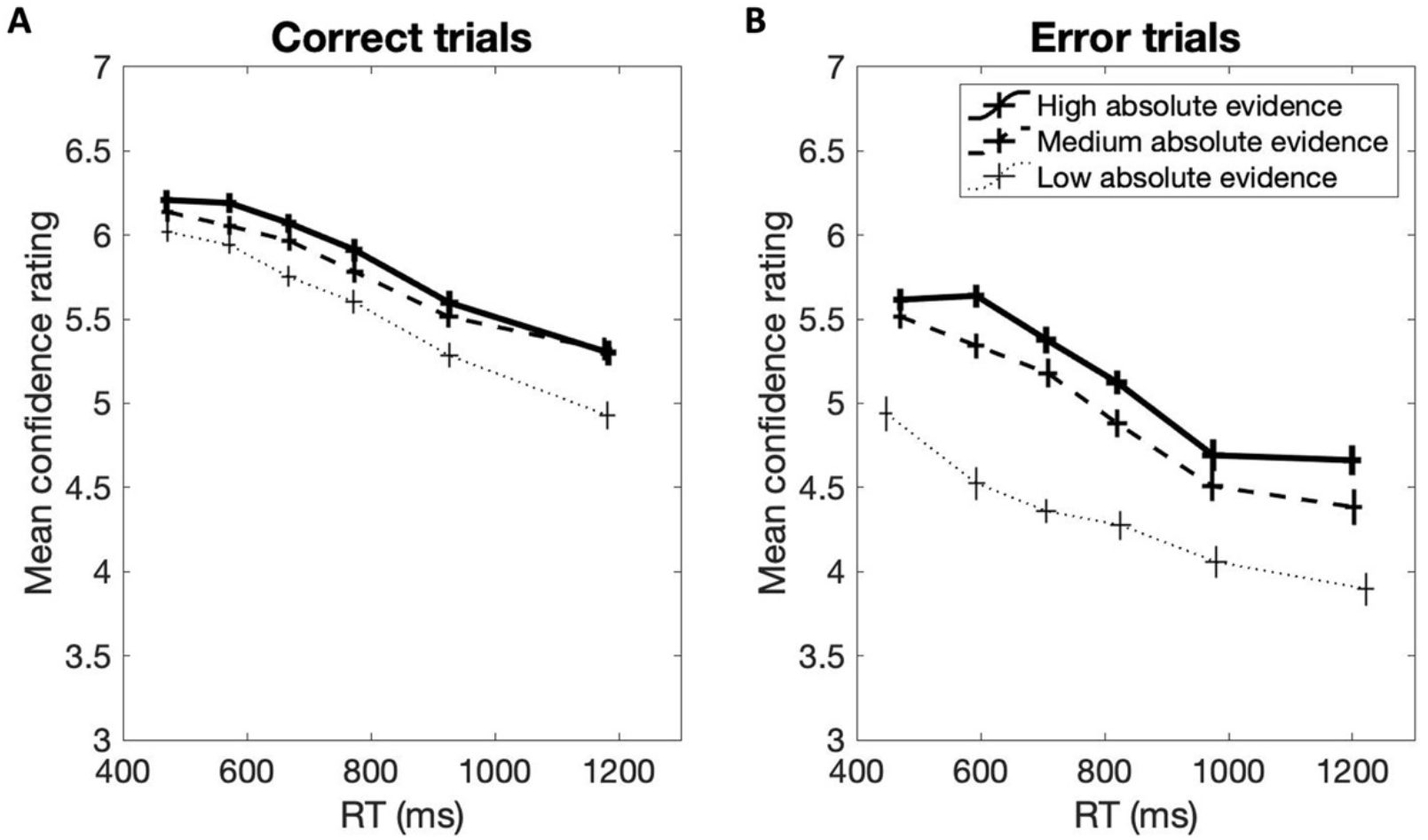
Experiment 1 mean RTs and confidence ratings by absolute evidence level and RT quantile bin (with borders between bins at the 10^th^, 30^th^, 50^th^, 70^th^, and 90^th^ RT percentiles) for (A) correct trials and (B) error trials. Horizontal and vertical error bars indicate SEM of RT and confidence, respectively.

In summary, these results confirm that, even though decision accuracy decreased with increasing absolute evidence, confidence increased for both correct and incorrect responses. Changes of mind likelihood results showed the opposite pattern, which could be expected from its negative relationship with confidence. Lastly, the positive effect of absolute evidence on confidence did not appear to be simply due to faster RTs in conditions with higher absolute evidence. Next, we investigated in Experiment 2 whether confidence also co-varied with stimulus variability in our task, which was introduced as an additional factor.

## Experiment 2

### Methods

#### Participants

A different sample of 35 university student volunteers with normal or corrected-to-normal vision was recruited. Six participants were subsequently excluded: Three failed to report confidence in more than 20% of all trials, one showed lower than 55% accuracy, and two reported the same confidence level in more than 90% of trials where confidence was reported. Twenty-nine participants were included in the analysis (mean age = 23, SD = 5, range 18 – 39 years, 25 females). This experiment was approved by the University of Melbourne ethics committee (ID: 1954641.2).

#### Experimental Procedures

Experimental procedures were identical to Experiment 1, except where noted below.

#### Task and Stimuli

Experiment 2 aimed to test the effect of evidence variability (i.e., the standard deviations of the distributions from which luminance values were sampled in each frame) on decision accuracy and confidence. It was a replication of Experiment 1 but included the additional factor “luminance variability”. We again used three levels of absolute evidence (low, medium, high), as in Experiment 1. We only included two levels of relative evidence (low, high) because the effects of relative evidence were not of primary interest in this experiment. The mean luminance values for the low and high relative evidence conditions were in-between the values used in Experiment 1 (see Supplementary Table S19) to reduce ceiling and floor effects. We further included two levels of luminance variability (low, high), resulting in a 3 × 2 × 2 design. Evidence variability was operationalised as the variability of the luminance value distributions (standard deviation of 25.5 for high variability and 12.5 for low variability; Supplementary Figure S3). The high variability condition was identical to Experiment 1, while the low variability condition contained only half the variability around the mean as compared to Experiment 1. The task structure, stimulus presentation, and apparatus were otherwise the same as in Experiment 1.

#### Data Analysis

The same GLMM approach used in Experiment 1 was used for Experiment 2, except that the models included the additional factor of variability and its interactions with relative evidence, absolute evidence, and the three-way interaction term.

### Results

#### Accuracy and Response Times

For decision accuracy, there were an effect of relative evidence (*χ²*[1] = 576.22, *p* <.001), an effect of absolute evidence (*χ²*[2] = 588.29, *p* <.001), and an interaction between relative and absolute evidence (*χ²*[2] = 37.02, *p* <.001) as in Experiment 1. This interaction was again because log odds of being correct were reduced by absolute evidence more strongly when relative evidence was high (see Supplementary Figure S4). Importantly, luminance variability and its interaction terms were not significant. As depicted in Figures 6A and 6D, the accuracy of participants’ responses increased with stronger relative evidence and decreased with higher absolute evidence, regardless of flicker variability. Full statistical results are presented in Supplementary Tables S20 and S21.

**Figure 6.**
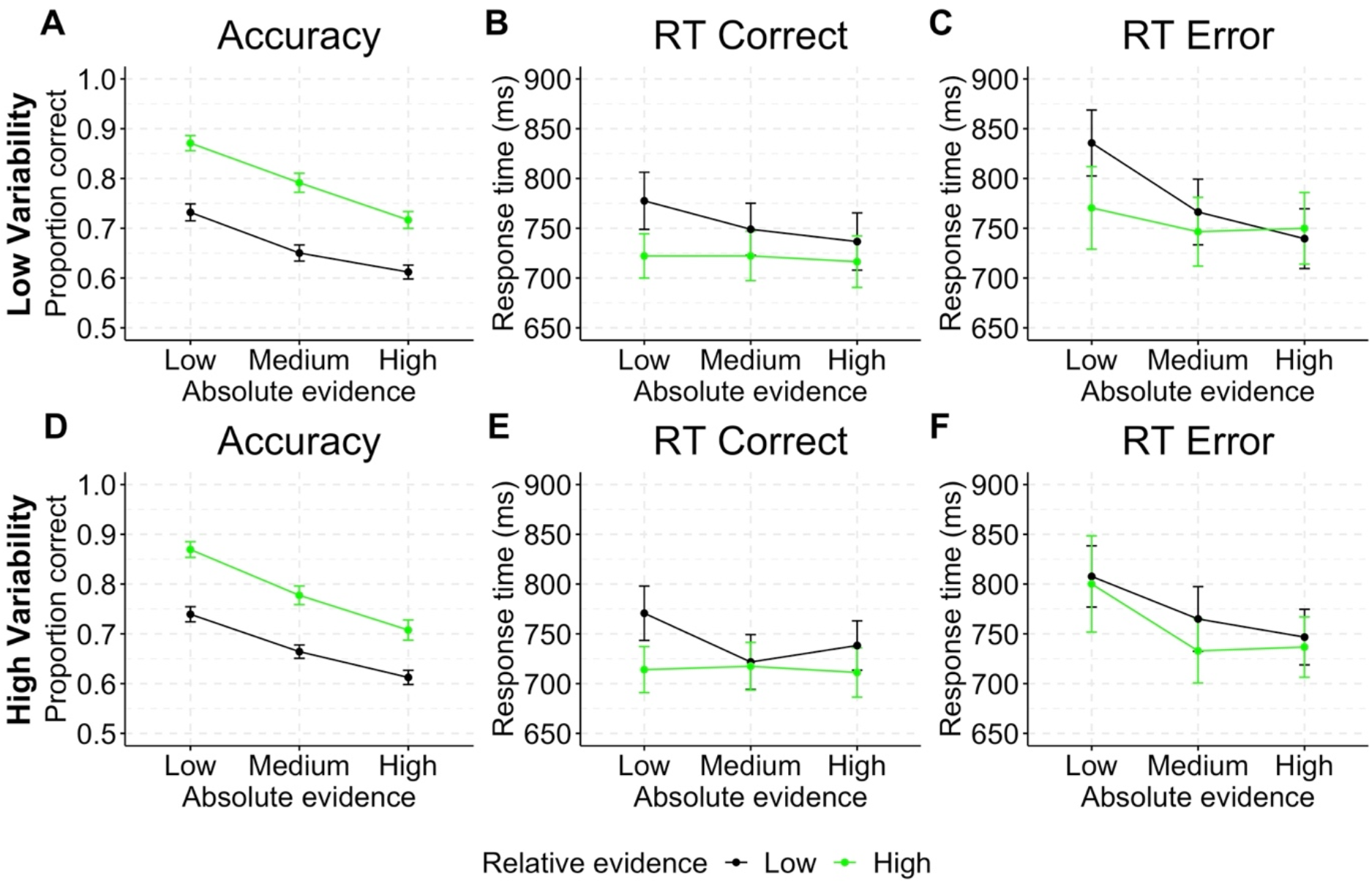
Experiment 2 accuracy and response time (RT). (A, D) Decision accuracy (average proportion correct) for the low (A) and high (D) luminance variability conditions. (B, E) Mean RTs for correct trials for low (B) and high (E) luminance variability conditions. (C, F) Mean RTs for error trials for low (C) and high (F) luminance variability conditions. Error bars represent SEM.

For RTs, when analysing data from correct trials, there was an effect of relative evidence (*χ²*[1] = 106.53, *p* <.001), an effect of absolute evidence (*χ²*[2] = 44.46, *p* <.001), and an interaction between relative and absolute evidence (*χ²*[2] = 32.20, *p* <.001), again replicating Experiment 1 (Figures 6B and 6E). Additionally, we found an effect of luminance variability (*χ²*[1] = 6.61, *p* = .010), an interaction between absolute evidence and luminance variability (*χ²*[2] = 7.13, *p* = .028), and an interaction among all three factors (*χ²*[2] = 6.24, *p* = .044). This effect appears to be driven by a small dip in RTs for the low relative / medium absolute evidence condition in the high compared to the low luminance variability condition. The overall pattern of RT results, however, was highly similar between luminance variability conditions, suggesting no substantial effect of luminance variability on RT in correct trials.

Analyses of error trials again showed a similar pattern of results (relative evidence: *χ²*[1] = 23.74, *p* <.001; absolute evidence: *χ²*[2] = 37.22, *p* <.001; interaction between relative and absolute evidence: *χ²*[2] = 14.25, *p* <.001), and no effect of luminance variability or interaction involving luminance variability. Figures 6C and 6F show that the RT effects for error trials from Experiment 1 replicated regardless of variability condition. Full statistical results are presented in Supplementary Tables S22 – S25.

Taken together, the effects of relative and absolute evidence on accuracy and RT found in Experiment 1 were replicated in Experiment 2. Importantly, we did not find strong and significant effects of luminance variability on accuracy or RT measures.

#### Confidence

When analysing correct trials, there was an effect of relative evidence (*χ²*[1] = 261.57, *p* <.001), an effect of absolute evidence (*χ²*[2] = 191.90, *p* <.001), and an interaction between relative and absolute evidence (*χ²*[2] = 47.83, *p* <.001). Confidence was higher in trials with stronger relative evidence and higher absolute evidence, and the effect of relative evidence was diminished in high absolute evidence conditions, replicating the pattern of results of Experiment 1. There was no significant effect of luminance variability nor significant interactions with luminance variability (Figures 7A and 7C).

**Figure 7.**
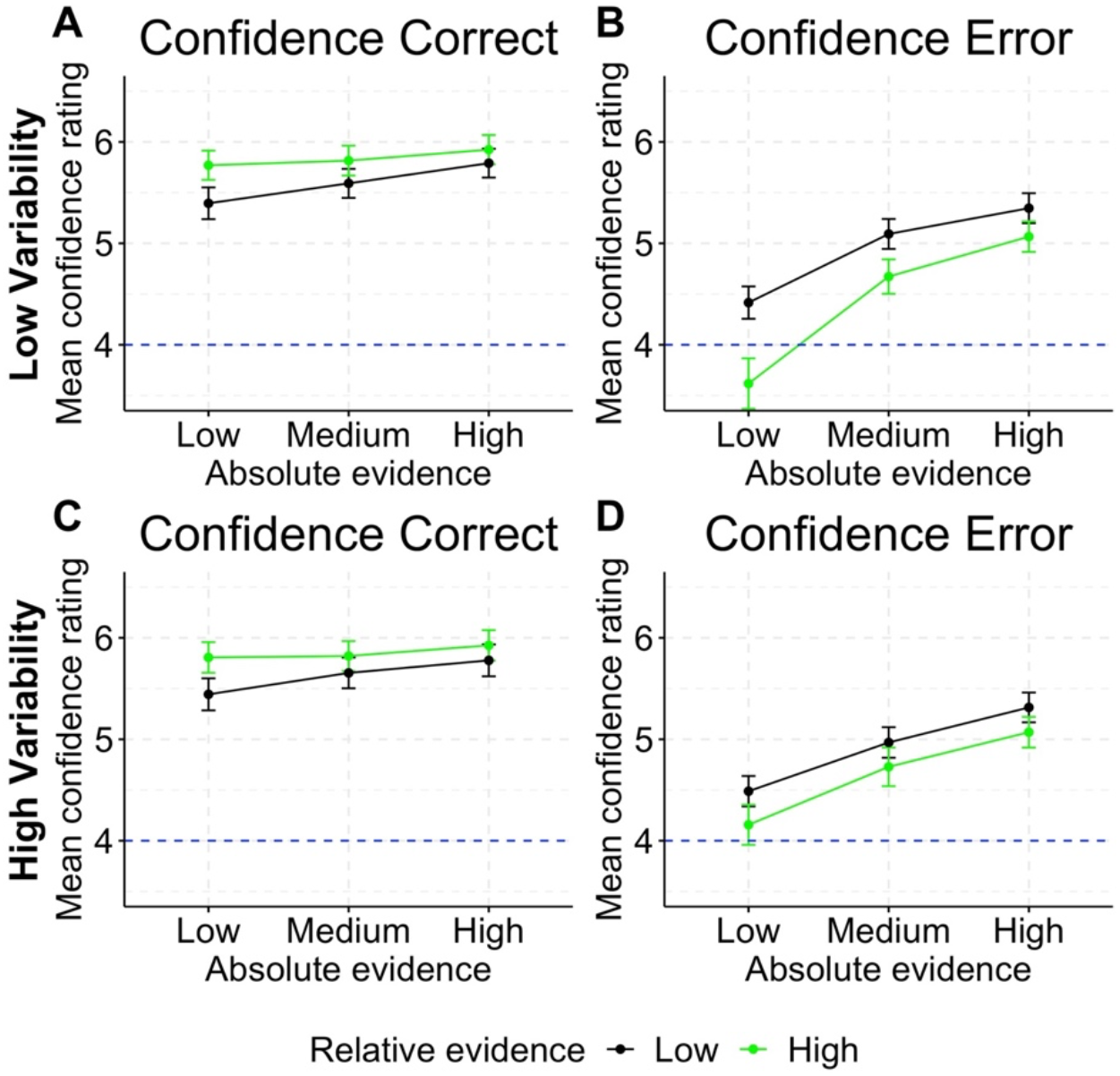
Mean confidence ratings in Experiment 2. (A, C) Mean confidence for correct trials for low (A) and high (C) variability. (B, D) Mean confidence for error trials for low (B) and high (D) variability. Confidence ratings were measured on a scale ranging from 1 (“surely incorrect”) to 7 (surely correct). The dotted line indicates the mid-point of the scale. Error bars represent SEM.

For analyses of error trials, we again found effects for relative and absolute evidence, but unlike in Experiment 1, despite reproducing the same overall pattern, their interaction failed to reach significance (relative evidence: *χ²*[1] =57.71, *p* <.001; absolute evidence: *χ²*[2] = 400.02, *p* <.001; Figures 7B and 7D). The absence of this interaction is most likely because we included only two levels of relative evidence, which were more similar to each other than in Experiment 1. While there was no main effect of luminance variability, an interaction was found between relative evidence and luminance variability (*χ²*[2] = 4.35, *p* =.037); as well as between absolute evidence and luminance variability (*χ²*[2] = 8.47, *p* = .014). The interaction between relative evidence and luminance variability was because higher variability was associated with increased confidence when relative evidence was high. That is, when relative evidence was high, low luminance variability was associated with lower confidence judgements, despite no detected change in performance (see above). The interaction between absolute evidence and luminance variability was reflected in higher luminance variability associated with increased confidence when absolute evidence was low. Full statistical results are presented in Supplementary Tables S26 – S29.

#### Change of Mind

For change-of-mind likelihood in correct trials, there was a negative effect of relative evidence (*χ²*[2] = 36.33, *p* <.001), and an interaction between relative and absolute evidence (*χ²*[2] = 9.72, *p* = .008). However, unlike in Experiment 1, there was no effect of absolute evidence (Figure 8A and 8C). For error trials, there was a positive effect of relative evidence (*χ²*[2] = 119.24, *p* <.001), a negative effect of absolute evidence (*χ²*[2] = 19.63, *p* <.001) and an interaction between relative and absolute evidence (*χ²*[2] = 125.93, *p* <.001; Figures 8B and 8D), as in Experiment 1. Full statistical results are presented in Supplementary Tables S30 – S33.

**Figure 8.**
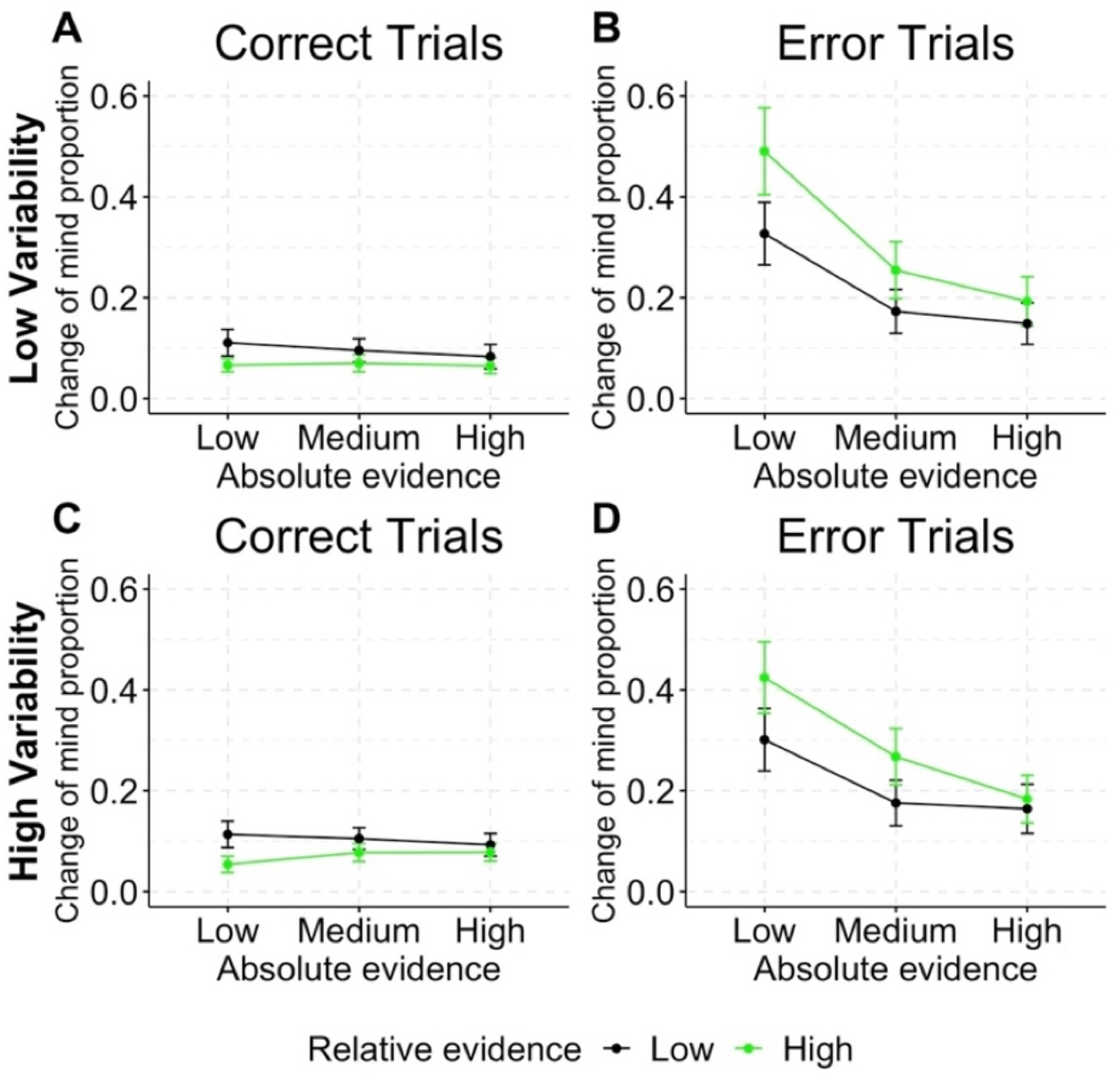
Experiment 2 proportions of change-of-mind trials in each condition. (A, C) Mean proportion of change of mind for correct trials for low (A) and high (C) variability. (B, D) Mean proportion of change of mind for error trials for low (B) and high (D) variability. Change-of-mind trials were defined as trials with confidence ratings lower than 4, indicating that the participant believed their brightness judgement was incorrect.

#### The Effect of Absolute Evidence on Confidence in Addition to RT

When modelling confidence in correct response trials using predictors of RT, luminance variability and relative evidence (but not absolute evidence), we observed both the effects of relative evidence (*χ²*[1] = 195.23, *p* <.001) and RT (*χ²*[1] = 1044.45, *p* <.001). When we added absolute evidence (and interactions involving this variable) as a predictor, we also observed the effect of absolute evidence (*χ²*[2] = 169.31, *p* <.001), and an interaction between absolute evidence and relative evidence (*χ²*[2] = 33.46, *p* <.001). For error trials, confidence was predicted by relative evidence (*χ²*[1] = 75.31, *p* <.001) and RT (*χ²*[1] = 357.34, *p* <.001). Additionally, it was predicted by absolute evidence (*χ²*[2] = 357.69, *p* <.001), an interaction between luminance variability and absolute evidence (*χ²*[2] = 6.83, *p* = .033), and an interaction between luminance variability and relative evidence (*χ²*[1] = 4.95, *p* = .026). Consistent with Experiment 1, Figures 9A and 9B also showed that absolute evidence increased confidence across RT bins for both correct and error trials. Full statistical results are presented in Supplementary Tables S34 and S37.

**Figure 9.**
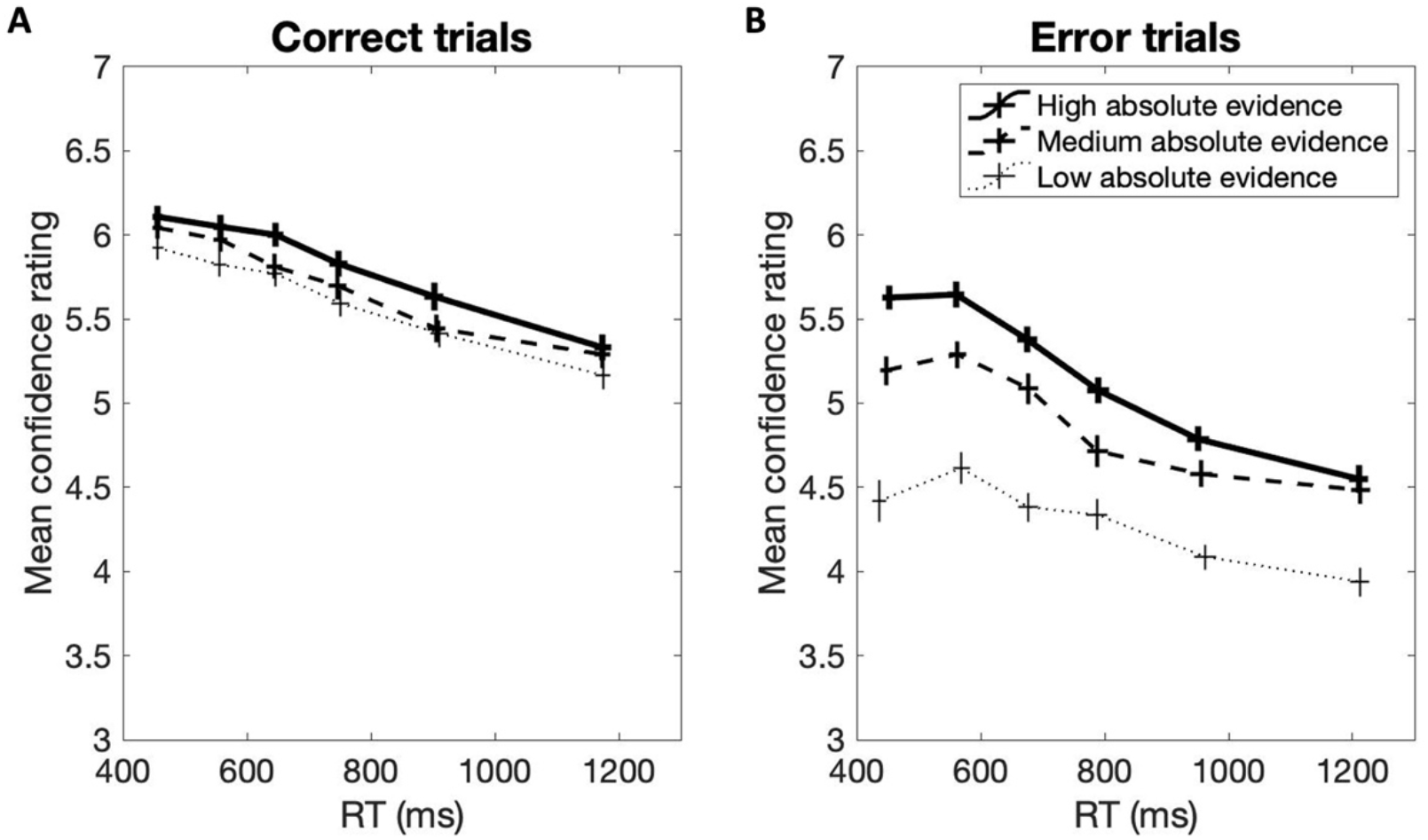
Experiment 2 mean RTs and confidence ratings by absolute evidence level and RT quantile bin (with borders between bins at the 10^th^, 30^th^, 50^th^, 70^th^, and 90^th^ RT percentiles) for (A) correct trials and (B) error trials. Horizontal and vertical error bars indicate SEM of RT and confidence, respectively.

Taken together, the results of Experiment 2 replicated the effects of relative and absolute evidence on decision accuracy and RT from Experiment 1. They further replicated the overall patterns of results for effects of relative and absolute evidence on confidence. The result that increasing absolute evidence led to increased confidence and faster RTs was also replicated. However, the effects on change-of-mind frequency were only replicated for error trials, but not for correct trials. This is possibly due to the fact that the effect on change of mind was rather weak as participants rarely change their mind after a correct response, and in Experiment 2 the luminance values of the low relative evidence level were higher than that of Experiment 1, in which the effect was stronger.

The luminance variability manipulation did not appear to substantially affect decision performance or confidence judgements. Only when looking at error trials specifically we observed some interactions between luminance variability and confidence. Overall, the pattern of results for high variability looked highly similar to Experiment 1. However, firstly, with lower luminance variability, confidence for error trials in high relative evidence trials was reduced. Given that these were trials in which errors were indeed committed, this means the combination of low luminance variability and strong relative evidence made it easier for participants to sense that their decision might have been wrong. Secondly, the combination of low variability and low absolute evidence also led to decreased confidence in error responses. This again indicates that participants found it somewhat easier to sense that their decision might have been wrong. However, we do not interpret the reported interaction effect as strong evidence that variability substantially influenced confidence judgments in our designs. This is because these interaction effects were rather small compared with the effects of relative and absolute evidence, and they were only observed within low ranges of relative and absolute evidence.

## Discussion

Based on the previous finding that higher levels of absolute evidence were associated with less accurate perceptual decisions and slower changes of mind (Turner, Feuerriegel, et al., 2021), and suggestions that confidence in one’s decision might moderate the speed and likelihood of later changes of mind (van den Berg et al., 2016; Turner, Feuerriegel, et al., 2021), we asked whether increases in absolute evidence are associated with higher decision confidence. We used a luminance discrimination task that was highly similar to that in Turner, Feuerriegel, and colleagues (2021) and examined the effect of absolute evidence on decision accuracy, RTs and confidence ratings. In this task, to manipulate absolute evidence we varied the overall (summed) luminance across the two stimuli, in addition to manipulating relative evidence (i.e., their luminance difference). Experiment 1 first replicated previous findings showing that increases in absolute evidence are associated with decreased accuracy and faster RTs (Ratcliff et al., 2018; Teodorescu et al., 2016; Turner, Feuerriegel, et al., 2021). We also found that while stronger relative evidence increased confidence for correct trials and reduced confidence for error trials, absolute evidence increased confidence for both correct and error trials. We replicated these effects in Experiment 2 and did not identify any substantial effects of luminance variability manipulations on task performance or decision confidence. Our findings suggest that heuristic biases in decision confidence associated with absolute evidence magnitude may ultimately make decisions harder to be subsequently overruled, in line with recent theoretical accounts (van den Berg et al., 2016; Turner, Feuerriegel et al., 2021).

### Why does high absolute evidence lead to decreased decision accuracy but increased confidence?

Increasing absolute evidence led to reduced accuracy but increased confidence. While seemingly paradoxical, these divergent effects have several coherent explanations within the general framework of an evidence accumulation process.

Considering first the negative effect of absolute evidence on decision accuracy, this can be explained, at least in part, by Weber’s law (Geisler, 1989; Ratcliff et al., 2018; Teodorescu et al., 2016; Turner, Feuerriegel, et al., 2021). Weber’s law suggests that relative evidence is perceptually reduced when absolute evidence is increased, due to nonlinear compressive scaling of the incoming sensory information. Over and above the effects of this compressive scaling, however, increases in absolute evidence are also thought to increase noise within the evidence accumulation process (Ratcliff et al., 2018; Turner, Feuerriegel et al., 2021). This could be explained by assuming that the variability of brightness representations scales with their mean luminance, such that more intense (i.e., brighter) stimuli are represented more variably in terms of brightness (Ratcliff et al., 2018). By assuming an input-dependent increase in noise within the decision process it is possible to account for both the speed up in initial RT and the decrease in choice consistency, which have been observed in previous studies under conditions of high absolute evidence (Ratcliff et al., 2018; Turner, Feuerriegel et al., 2021).

Turning now to consider the effect of absolute evidence magnitude on decision confidence, we found that increasing absolute evidence increased confidence, for both correct and incorrect decisions, despite the simultaneous decrease in decision accuracy. Hereafter, we will consider two possible explanations for this effect.

Firstly, this finding is in line with more recent studies that showed a positive evidence bias for decision confidence (Koizumi et al., 2015; Odegaard et al., 2018; Samaha et al., 2016; Samaha & Denison, 2020). In these studies, confidence was increased by experimentally increasing positive evidence (i.e., the extent of sensory evidence for the chosen decision outcome) while maintaining the ratio between positive and negative evidence. In other words, confidence judgments appeared to be based on the absolute magnitude of decision-congruent evidence but not decision-incongruent evidence.

These observations have led to development of several models of the processes that underlie decisions and confidence judgements (Maniscalco et al., 2021; Peters et al., 2017; Zylberberg et al., 2012). Central to all these models is the assumption that decisions and confidence judgements are based on two distinct sources of sensory evidence. While perceptual decisions are determined by relative evidence, confidence involves the heuristic use of only decision-congruent evidence.

From this viewpoint, the divergence between confidence and accuracy which we observed can be simply explained. For the decision process, relative evidence drove the decision outcome, with higher levels of absolute evidence leading to a decreased signal (due to Weber-scaling) and increased variability (due to input-dependent noise), within the decision process. As a result, decisions were, on average, faster and less accurate. Coincidently, due to our absolute evidence manipulation, the amount of decision-congruent evidence was boosted in conditions of high absolute evidence, leading to an increase in confidence, irrespective of these co-occurring accuracy and RT effects. In other words, the combined effects of Weber-scaling and a positive evidence bias can account for the effects of absolute evidence on decision accuracy, RTs, and confidence.

An alternative explanation for our divergent accuracy and confidence effects is that the increase in confidence we observed in high absolute evidence trials may have been a by-product of faster RTs. Certain models, distinct from those discussed above, suggest that confidence is partly determined by the time taken to come to a decision (e.g., Kiani et al., 2014; Zylberberg et al., 2016). As faster decision times are often associated with more reliable sources of evidence and correct decisions, RTs may inform confidence judgments. Considering the current findings, this view suggests that the increase we observed in decision confidence following increases in absolute evidence might have been due to the co-occurring speed up in response times. However, our results showed that, while higher absolute evidence led to both faster RTs and increased confidence, and faster RTs co-occurred with higher confidence ratings, RT alone could not fully explain the effect of absolute evidence on confidence. Within similar RT ranges, increased absolute evidence was still associated with increased confidence. This provides evidence that the effect of absolute evidence on confidence was not simply due to speeding of RTs.

### How does the effect of absolute evidence on confidence translate to change-of-mind decisions?

Our findings show that increasing absolute evidence magnitude, which led to slower change-of-mind decisions in an almost identical design (Turner, Feuerriegel, et al., 2021), also increases participants’ sense of confidence in their decisions. Moreover, when confidence ratings were coded as a binary change-of-mind variable, increasing absolute evidence similarly led to reduced change-of-mind frequency (except for Experiment 2, correct trials). This supports that confidence and changes of mind in our design were indeed closely related and both depended on absolute evidence. This finding is consistent with the idea that subjective feelings of confidence in one’s decision may affect subsequent change-of-mind decisions (van den Berg et al., 2016). More generally, this implies that heuristic biases in confidence judgements, such as those associated with absolute evidence magnitude/the positive evidence bias (e.g., Peters et al., 2017; Zylberberg et al., 2012), or motor-related confidence biases (e.g., Fleming & Daw, 2017; Turner, Angdias, et al., 2021; Gajdos et al., 2019) may play an essential role in determining the speed and likelihood of subsequent change-of-mind decisions.

The positive association we found between absolute evidence magnitude and decision confidence may be important to consider when attempting to model the dynamics of error correction and changes of mind. For example, it was recently shown that existing change-of-mind models, based solely on the ongoing accumulation of relative evidence after a decision, cannot easily account for the effects of absolute evidence on change-of-mind likelihood or RTs (Turner, Feuerriegel, et al., 2021). In particular, these models have difficulty capturing patterns of slower change-of-mind RTs in higher absolute evidence conditions. To completely account for decision and changes-of-mind behaviour, novel modelling assumptions may need to be explored. For example, one possibility is that the decision threshold for changing one’s mind may depend, at least in part, on decision confidence. That is, decisions made with higher confidence may require more significant amounts of contradictory evidence to trigger a decision reversal (Turner, Feuerriegel, et al., 2021; see also van den Berg et al., 2016). Alternatively, it is possible that the weighting of post-decisional evidence may depend on decision confidence (Braun et al., 2018; Rollwage et al., 2020). In other words, the dynamics of post-decisional processing may be fundamentally altered in a confidence-depended manner.

By demonstrating that decision confidence does vary across changes in absolute evidence magnitude, the current study provides empirical backing for exploring such assumptions. Suppose future theoretical work were to incorporate confidence-related biases in specific model parameters (such as shifts in the change-of-mind threshold) within existing computational frameworks, this may yield better accounts of the dynamics underlying change-of-mind decisions, and may help to integrate insights from recent theoretical accounts that capture effects of absolute evidence and positive evidence biases (e.g., Maniscalco et al., 2021).

### Limitations

Our findings should be interpreted with the following caveats in mind. Firstly, one difference between our study and Turner, Feuerriegel, and colleagues’ (2021) study that limits the generalizability of our confidence findings to their change-of-mind results is that in their study, stimuli remained on the screen for a short duration after the initial response was submitted, while in our study the stimuli were terminated after the response. It should be noted that it is reasonable to assume that visual processing continues for around 300 ms after a stimulus is terminated (Resulaj et al., 2009), reducing the relevance of this issue. However, it is still possible that the difference in trial structure prompted temporally different computations. Change-of-mind decisions in Turner, Feuerriegel, and colleagues (2021) might have been more strongly based on late-arriving post-decisional evidence (e.g., Charles & Yeung, 2019), while our confidence judgements could not be based on such information. Future studies could investigate whether such different presentation conditions might prompt participants to use incoming sensory evidence differently to compute confidence. If a change-of-mind is the end-product of a confidence computation, this could affect how confidence is reported.

Another limitation is that we did not include an extensive range of luminance variability conditions in Experiment 2. This means that we cannot rule out the possibility that more extensive magnitude manipulations of luminance variability might exert more substantial effects on task performance and confidence measures in our task. For example, there are many examples of stimulus variability-related effects in other decision tasks (e.g., Desender et al., 2018; Zylberberg et al., 2014), although these are not always consistent in the direction of their effects. However, given that the observed effects appeared to be very small in our study, we believe that changes in perceived variability are an unlikely explanation for the much larger and consistent effects of absolute evidence in our experiments.

## Conclusion

Using a luminance discrimination task, we showed that stronger absolute evidence led to reduced decision accuracy, faster RTs, and increased decision confidence. This finding parallels previous findings of higher absolute evidence leading to slower changes of mind (Turner, Feuerriegel et al., 2021) and suggests that decision confidence may moderate the speed of decision reversals in perceptual judgment tasks. Our results are compatible with recent suggestions that confidence might be driven by decision-congruent evidence, in addition to the theory that faster response time contributes to higher confidence. Finally, by demonstrating that decision confidence varies with absolute evidence magnitude, the current work provides an empirical basis for future exploration of potential confidence-related changes in post-decisional information processing (e.g., shifts in the change-of-mind threshold or changes in the weighting of evidence).

## Acknowledgements

The authors thank all colleagues from the Decision Neuroscience Lab, and the Institute of Neuroscience and Medicine (INM-3), Cognitive Neuroscience, for discussions and support, and Megan Peters and Brian Maniscalco for valuable discussions. This work was funded by an Australian Research Council Discovery Project Grant (DP160103353) to S.B. and R.H. Y.H.K. and H.O. were funded by a Jülich University of Melbourne Postgraduate Academy (JUMPA) scholarship. G.R.F. and P.H.W. were funded by the Deutsche Forschungsgemeinschaft (DFG, German Research Foundation) – Project-ID 431549029 – SFB 1451.

