## Supplementary Material for "Divergent effects of absolute evidence magnitude on decision accuracy and confidence in perceptual judgements"

Table S9 Experiment 1 Likelihood Ratio Tests Results for Predicting Confidence (Error Trials) from Relative Evidence, Absolute Evidence, and Their Interactions ... 13

Table S10 Experiment 1 Regression Coefficients for Predicting Confidence (Error Trials) from Relative Evidence, Absolute Evidence, and Their Interactions ... 13

Table S11 Experiment 1 Likelihood Ratio Tests Results for Predicting Change of Mind (Log Odds of Confidence Lower Than 4) in Correct Trials from Relative Evidence, Absolute Evidence, and Their Interactions ... 14

Table S12 Experiment 1 Regression Coefficients for Predicting Change of Mind (Log Odds of Confidence Lower Than 4) in Correct Trials from Relative Evidence, Absolute Evidence, and Their Interactions ... 14

Table S13 Experiment 1 Likelihood Ratio Tests Results for Predicting Change of Mind (Log Odds of Confidence Lower Than 4) in Error Trials from Relative Evidence, Absolute Evidence, and Their Interactions ... 15

Table S14 Experiment 1 Regression Coefficients for Predicting Change of Mind (Log Odds of Confidence Lower Than 4) in Error Trials from Relative Evidence, Absolute Evidence, and Their Interactions ... 15

Table S15 Experiment 1 Likelihood Ratio Tests Results for Predicting Confidence (Correct Trials) from Relative Evidence, RT, Absolute Evidence, and Their Interactions ... 16

Table S16 Experiment 1 Regression Coefficients for Predicting Confidence (Correct Trials) from Relative Evidence, RT, Absolute Evidence, and Their Interactions ... 17

Table S17 Experiment 1 Likelihood Ratio Tests Results for Predicting Confidence (Error Trials) from Relative Evidence, RT, Absolute Evidence, and Their Interactions ... 18

Table S18 Experiment 1 Regression Coefficients for Predicting Confidence (Error Trials) from Relative Evidence, RT, Absolute Evidence, and Their Interactions ........ 19

Figure S3. Example of luminance value distributions. ........ 20

Figure S4. Experiment 2 mean log odds of being correct in each condition ........ 21

Table S19 Mean luminance values for all experimental conditions of Experiment 2 ........ 22

Table S20 Experiment 2 Likelihood Ratio Tests Results for Predicting Accuracy (Log Odds of Being Correct) from Relative Evidence, Absolute Evidence, Luminance Variability, and Their Interactions ........ 23

Table S21 Experiment 2 Regression Coefficients for Predicting Accuracy (Log Odds of Being Correct) from Relative Evidence, Absolute Evidence, Luminance Variability, and Their Interactions ........ 24

Table S22 Experiment 2 Likelihood Ratio Tests Results for Predicting Response Time (Correct Trials) from Relative Evidence, Absolute Evidence, Luminance Variability, and Their Interactions ........ 25

Table S23 Experiment 2 Regression Coefficients for Predicting Response Time (Correct Trials) from Relative Evidence, Absolute Evidence, Luminance Variability, and Their Interactions ........ 26

Table S24 Experiment 2 Likelihood Ratio Tests Results for Predicting Response Time (Error Trials) from Relative Evidence, Absolute Evidence, Luminance Variability, and Their Interactions ........ 27

Table S25 Experiment 2 Regression Coefficients for Predicting Response Time (Error Trials) from Relative Evidence, Absolute Evidence, Luminance Variability, and Their Interactions 28

Table S26 Experiment 2 Likelihood Ratio Tests Results for Predicting Confidence (Correct Trials) from Relative Evidence, Absolute Evidence, Luminance Variability, and Their Interactions ..... 29

Table S27 Experiment 2 Regression Coefficients for Predicting Confidence (Correct Trials) from Relative Evidence, Absolute Evidence, Luminance Variability, and Their Interactions 30

Table S28 Experiment 2 Likelihood Ratio Tests Results for Predicting Confidence (Error Trials) from Relative Evidence, Absolute Evidence, Luminance Variability, and Their Interactions ..... 31

Table S29 Experiment 2 Regression Coefficients for Predicting Confidence (Error Trials) from Relative Evidence, Absolute Evidence, Luminance Variability, and Their Interactions 32

Table S30 Experiment 2 Likelihood Ratio Tests Results for Predicting Change of Mind (Log Odds of Confidence Lower Than 4) in Correct Trials from Relative Evidence, Absolute Evidence, Luminance Variability, and Their Interactions ..... 33

Table S31 Experiment 2 Regression Coefficients for Predicting Change of Mind (Log Odds of Confidence Lower Than 4) in Correct Trials from Relative Evidence, Absolute Evidence, Luminance Variability, and Their Interactions ..... 34

Table S32 Experiment 2 Likelihood Ratio Tests Results for Predicting Change of Mind (Log Odds of Confidence Lower Than 4) in Error Trials from Relative Evidence, Absolute Evidence, Luminance Variability, and Their Interactions ..... 35

Table S33 Experiment 2 Regression Coefficients for Predicting Change of Mind (Log Odds of Confidence Lower Than 4) in Error Trials from Relative Evidence, Absolute Evidence, Luminance Variability, and Their Interactions ..... 36

Table S34 Experiment 2 Likelihood Ratio Tests Results for Predicting Confidence (Correct Trials) from Relative Evidence, Luminance Variability, RT, Absolute Evidence, and Their Interactions ..... 37

Table S35 Experiment 2 Regression Coefficients for Predicting Confidence (Correct Trials) from Relative Evidence, Luminance Variability, RT, Absolute Evidence, and Their Interactions ..... 38

Table S36 Experiment 2 Likelihood Ratio Tests Results for Predicting Confidence (Error Trials) from Relative Evidence, Luminance Variability, RT, Absolute Evidence, and Their Interactions ..... 39

Table S37 Experiment 2 Regression Coefficients for Predicting Confidence (Error Trials) from Relative Evidence, Luminance Variability, RT, Absolute Evidence, and Their Interactions ..... 40

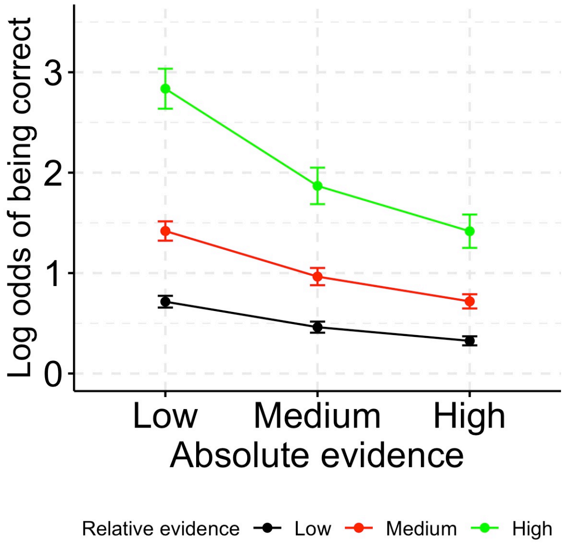

Figure S1. Experiment 1 mean log odds of being correct in each condition. Error bars represent SEM.

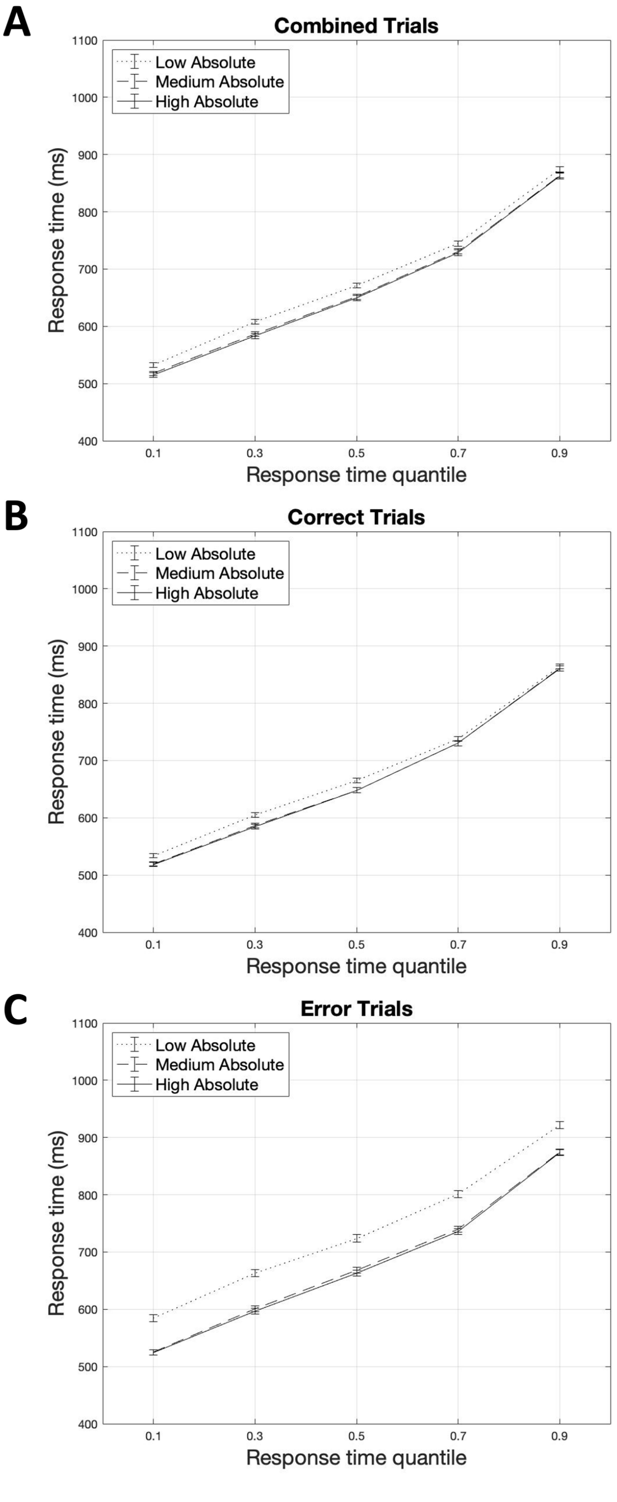

Figure S2. Experiment 1 response time quantiles across absolute evidence levels (collapsing across relative evidence levels). (A) Correct and error trials combined. (B) Correct trials. (C) Error trials. Error bars represent SEM.

**Accuracy (Log Odds of Being Correct)**

Table S1
*Experiment 1 Likelihood Ratio Tests Results for Predicting Accuracy (Log Odds of Being Correct) from Relative Evidence, Absolute Evidence, and Their Interactions*

| Predictor | *df* | *χ2* | *p* |
|---|---|---|---|
| Rel | 2 | 1433.01 | <.001*** |
| Abs | 2 | 485.87 | <.001*** |
| Rel × Abs | 4 | 91.71 | <.001*** |

Note. Rel: Relative evidence; Abs: Absolute evidence.
*p <.05 **p <.01 ***p <.001

Table S2
*Experiment 1 Regression Coefficients for Predicting Accuracy (Log Odds of Being Correct) from Relative Evidence, Absolute Evidence, and Their Interactions*

| Parameters | Estimate | *SE* | *z* | *p* |
|---|---|---|---|---|
| Intercept | 1.11 | 0.07 | 16.47 | <.001*** |
| Low Rel | -0.60 | 0.02 | -32.09 | <.001*** |
| Med Rel | -0.10 | 0.02 | -4.97 | <.001*** |
| Low Abs | 0.43 | 0.02 | 19.56 | <.001*** |
| Med Abs | -0.09 | 0.02 | -4.50 | <.001*** |
| Low Rel × Low Abs | -0.22 | 0.03 | -7.72 | <.001*** |
| Med Rel × Low Abs | -0.06 | 0.03 | -2.16 | .031* |
| Low Rel × Med Abs | 0.05 | 0.03 | 2.04 | .041* |
| Med Rel × Med Abs | 0.02 | 0.03 | 0.92 | .356 |

Note. Intercept represents the estimate for high relative and high absolute evidence. Rel: Relative evidence; Abs: Absolute evidence.
*p <.05 **p <.01 ***p <.001

**Response Time (Correct Trials)**

Table S3
*Experiment 1 Likelihood Ratio Tests Results for Predicting Response Time (Correct Trials) from Relative Evidence, Absolute Evidence, and Their Interactions*

| Predictor | *df* | $\chi 2$ | *p* |
|---|---|---|---|
| Rel | 2 | 314.71 | <.001*** |
| Abs | 2 | 35.85 | <.001*** |
| Rel × Abs | 4 | 106.25 | <.001*** |

Note. Rel: Relative evidence; Abs: Absolute evidence.
*p <.05 **p <.01 ***p <.001

Table S4
*Experiment 1 Regression Coefficients for Predicting Response Time (Correct Trials) from Relative Evidence, Absolute Evidence, and Their Interactions*

| Parameters | Estimate | *SE* | *z* | *p* |
|---|---|---|---|---|
| Intercept | 765.12 | 2.84 | 269.20 | <.001*** |
| Low Rel | 24.69 | 1.42 | 17.43 | <.001*** |
| Med Rel | 6.31 | 1.14 | 5.55 | <.001*** |
| Low Abs | 11.16 | 1.53 | 7.31 | <.001*** |
| Med Abs | -6.39 | 1.32 | -4.83 | <.001*** |
| Low Rel × Low Abs | 19.47 | 1.89 | 10.31 | <.001*** |
| Med Rel × Low Abs | 4.06 | 1.30 | 3.12 | .002** |
| Low Rel × Med Abs | -1.88 | 1.45 | -1.29 | .195 |
| Med Rel × Med Abs | -4.33 | 1.66 | -2.61 | .009** |

Note. Intercept represents the estimate for high relative and high absolute evidence. Rel: Relative evidence; Abs: Absolute evidence.
*p <.05 **p <.01 ***p <.001

**Response Time (Error Trials)**

Table S5

*Experiment 1 Likelihood Ratio Tests Results for Predicting Response Time (Error Trials) from Relative Evidence, Absolute Evidence, and Their Interactions*

| Predictor | *df* | $\chi 2$ | *p* |
|---|---|---|---|
| Rel | 2 | 18.54 | <.001*** |
| Abs | 2 | 49.96 | <.001*** |
| Rel × Abs | 4 | 20.20 | <.001*** |

Note. Rel: Relative evidence; Abs: Absolute evidence.
*p <.05 **p <.01 ***p <.001

Table S6

*Experiment 1 Regression Coefficients for Predicting Response Time (Error Trials) from Relative Evidence, Absolute Evidence, and Their Interactions*

| Parameters | Estimate | *SE* | *z* | *p* |
|---|---|---|---|---|
| Intercept | 811.14 | 7.68 | 105.63 | <.001*** |
| Low Rel | 12.87 | 2.62 | 4.92 | <.001*** |
| Med Rel | 2.11 | 2.67 | 0.79 | .427 |
| Low Abs | 24.84 | 2.75 | 9.02 | <.001*** |
| Med Abs | -5.51 | 2.86 | -1.93 | .054 |
| Low Rel × Low Abs | 21.36 | 3.70 | 5.77 | <.001*** |
| Med Rel × Low Abs | -10.33 | 3.76 | -2.75 | .006** |
| Low Rel × Med Abs | -14.42 | 3.59 | -4.02 | <.001*** |
| Med Rel × Med Abs | 7.49 | 3.78 | 1.98 | .048* |

Note. Intercept represents the estimate for high relative and high absolute evidence. Rel: Relative evidence; Abs: Absolute evidence.
*p <.05 **p <.01 ***p <.001

**Confidence (Correct Trials)**

Table S7

*Experiment 1 Likelihood Ratio Tests Results for Predicting Confidence (Correct Trials) from Relative Evidence, Absolute Evidence, and Their Interactions*

| Predictor | *df* | *χ2* | *p* |
|---|---|---|---|
| Rel | 2 | 879.07 | <.001*** |
| Abs | 2 | 293.89 | <.001*** |
| Rel × Abs | 4 | 121.55 | <.001*** |

Note. Rel: Relative evidence; Abs: Absolute evidence.
*p <.05 **p <.01 ***p <.001

Table S8

*Experiment 1 Regression Coefficients for Predicting Confidence (Correct Trials) from Relative Evidence, Absolute Evidence, and Their Interactions*

| Parameters | Estimate | *SE* | *z* | *p* |
|---|---|---|---|---|
| Intercept | 5.74 | 0.12 | 49.75 | <.001*** |
| Low Rel | -0.28 | 0.01 | -23.65 | <.001*** |
| Med Rel | -0.02 | 0.01 | -2.10 | .036* |
| Low Abs | -0.19 | 0.01 | -16.72 | <.001*** |
| Med Abs | 0.05 | 0.01 | 4.31 | <.001*** |
| Low Rel × Low Abs | -0.14 | 0.02 | -8.25 | <.001*** |
| Med Rel × Low Abs | -0.01 | 0.02 | -0.63 | .526 |
| Low Rel × Med Abs | 0.01 | 0.02 | 0.73 | .463 |
| Med Rel × Med Abs | 0.02 | 0.02 | 1.29 | .195 |

Note. Intercept represents the estimate for high relative and high absolute evidence. Rel: Relative evidence; Abs: Absolute evidence.
*p <.05 **p <.01 ***p <.001

**Confidence (Error Trials)**

Table S9

*Experiment 1 Likelihood Ratio Tests Results for Predicting Confidence (Error Trials) from Relative Evidence, Absolute Evidence, and Their Interactions*

| Predictor | *df* | $\chi 2$ | *p* |
|---|---|---|---|
| Rel | 2 | 99.99 | <.001*** |
| Abs | 2 | 392.15 | <.001*** |
| Rel × Abs | 4 | 9.30 | .054 |

Note. Rel: Relative evidence; Abs: Absolute evidence.
*p <.05 **p <.01 ***p <.001

Table S10

*Experiment 1 Regression Coefficients for Predicting Confidence (Error Trials) from Relative Evidence, Absolute Evidence, and Their Interactions*

| Parameters | Estimate | *SE* | *z* | *p* |
|---|---|---|---|---|
| Intercept | 4.77 | 0.13 | 35.48 | <.001*** |
| Low Rel | 0.25 | 0.03 | 9.93 | <.001*** |
| Med Rel | -0.02 | 0.03 | -0.70 | .486 |
| Low Abs | -0.58 | 0.03 | -18.52 | <.001*** |
| Med Abs | 0.14 | 0.03 | 5.23 | <.001*** |
| Low Rel × Low Abs | 0.01 | 0.04 | 0.18 | .855 |
| Med Rel × Low Abs | -0.02 | 0.04 | -0.41 | .681 |
| Low Rel × Med Abs | 0.08 | 0.03 | 2.34 | .019* |
| Med Rel × Med Abs | -0.02 | 0.04 | -0.63 | .531 |

Note. Intercept represents the estimate for high relative and high absolute evidence. Rel: Relative evidence; Abs: Absolute evidence.
*p <.05 **p <.01 ***p <.001

**Change of Mind (Log Odds of Confidence Lower Than 4) in Correct Trials**

Table S11

*Experiment 1 Likelihood Ratio Tests Results for Predicting Change of Mind (Log Odds of Confidence Lower Than 4) in Correct Trials from Relative Evidence, Absolute Evidence, and Their Interactions*

| Predictor | *df* | $\chi 2$ | *p* |
|---|---|---|---|
| Rel | 2 | 259.51 | <.001*** |
| Abs | 2 | 38.67 | <.001*** |
| Rel × Abs | 4 | 22.35 | <.001*** |

Note. Rel: Relative evidence; Abs: Absolute evidence.
*p <.05 **p <.01 ***p <.001

Table S12

*Experiment 1 Regression Coefficients for Predicting Change of Mind (Log Odds of Confidence Lower Than 4) in Correct Trials from Relative Evidence, Absolute Evidence, and Their Interactions*

| Parameters | Estimate | *SE* | *z* | *p* |
|---|---|---|---|---|
| Intercept | 3.58 | 0.29 | 12.44 | <.001*** |
| Low Rel | -0.53 | 0.04 | -12.81 | <.001*** |
| Med Rel | -0.14 | 0.04 | -3.31 | .001** |
| Low Abs | -0.26 | 0.04 | -6.31 | <.001*** |
| Med Abs | 0.13 | 0.04 | 3.02 | .003** |
| Low Rel × Low Abs | -0.17 | 0.05 | -3.19 | .001** |
| Med Rel × Low Abs | -0.08 | 0.06 | -1.40 | .162 |
| Low Rel × Med Abs | -0.04 | 0.06 | -0.60 | .550 |
| Med Rel × Med Abs | 0.07 | 0.06 | 1.14 | .255 |

Note. Intercept represents the estimate for high relative and high absolute evidence. Rel: Relative evidence; Abs: Absolute evidence.
*p <.05 **p <.01 ***p <.001

**Change of Mind (Log Odds of Confidence Lower Than 4) in Error Trials**

Table S13

*Experiment 1 Likelihood Ratio Tests Results for Predicting Change of Mind (Log Odds of Confidence Lower Than 4) in Error Trials from Relative Evidence, Absolute Evidence, and Their Interactions*

| Predictor | *df* | $\chi 2$ | *p* |
|---|---|---|---|
| Rel | 2 | 96.92 | <.001*** |
| Abs | 2 | 176.64 | <.001*** |
| Rel × Abs | 4 | 14.09 | .007** |

Note. Rel: Relative evidence; Abs: Absolute evidence.
*p <.05 **p <.01 ***p <.001

Table S14

*Experiment 1 Regression Coefficients for Predicting Change of Mind (Log Odds of Confidence Lower Than 4) in Error Trials from Relative Evidence, Absolute Evidence, and Their Interactions*

| Parameters | Estimate | *SE* | *z* | *p* |
|---|---|---|---|---|
| Intercept | 1.60 | 0.19 | 8.26 | <.001*** |
| Low Rel | 0.43 | 0.04 | 9.59 | <.001*** |
| Med Rel | -0.04 | 0.04 | -0.95 | .344 |
| Low Abs | -0.61 | 0.05 | -12.38 | <.001*** |
| Med Abs | 0.09 | 0.05 | 2.04 | .042* |
| Low Rel × Low Abs | 0.00 | 0.06 | 0.04 | .970 |
| Med Rel × Low Abs | -0.03 | 0.06 | -0.49 | .623 |
| Low Rel × Med Abs | 0.19 | 0.06 | 3.16 | .002** |
| Med Rel × Med Abs | -0.04 | 0.06 | -0.62 | .536 |

Note. Intercept represents the estimate for high relative and high absolute evidence. Rel: Relative evidence; Abs: Absolute evidence.
*p <.05 **p <.01 ***p <.001

**Confidence (Correct Trials)**

Table S15

*Experiment 1 Likelihood Ratio Tests Results for Predicting Confidence (Correct Trials) from Relative Evidence, RT, Absolute Evidence, and Their Interactions*

| Predictor | *df* | $\chi 2$ | *p* |
|---|---|---|---|
| Rel | 2 | 667.60 | <.001*** |
| RT.cmc | 1 | 1207.10 | <.001*** |
| Abs | 2 | 270.15 | <.001*** |
| Rel × Abs | 4 | 80.22 | <.001*** |

Note. Rel: Relative evidence; Abs: Absolute evidence;
RT.cmc: Response time (cluster-mean centered).
*p <.05 **p <.01 ***p <.001

Table S16

*Experiment 1 Regression Coefficients for Predicting Confidence (Correct Trials) from Relative Evidence, RT, Absolute Evidence, and Their Interactions*

| Parameters | Estimate | *SE* | *z* | *p* |
|---|---|---|---|---|
| Intercept | 5.75 | 0.12 | 49.77 | <.001*** |
| Low Rel | -0.24 | 0.01 | -20.99 | <.001*** |
| Med Rel | -0.01 | 0.01 | -1.27 | .205 |
| RT.cmc | -0.00 | 0.00 | -35.22 | <.001*** |
| Low Abs | -0.17 | 0.01 | -15.92 | <.001*** |
| Med Abs | 0.04 | 0.01 | 3.72 | <.001*** |
| Low Rel × Low Abs | -0.11 | 0.02 | -6.81 | <.001*** |
| Med Rel × Low Abs | -0.00 | 0.02 | -0.26 | .793 |
| Low Rel × Med Abs | 0.01 | 0.02 | 0.53 | .598 |
| Med Rel × Med Abs | 0.02 | 0.02 | 0.97 | .330 |

Note. Intercept represents the estimate for high relative and high absolute evidence. Rel: Relative evidence; Abs: Absolute evidence; RT.cmc: Response time (cluster-mean centered).
*p <.05 **p <.01 ***p <.001

**Confidence (Error Trials)**

Table S17

*Experiment 1 Likelihood Ratio Tests Results for Predicting Confidence (Error Trials) from Relative Evidence, RT, Absolute Evidence, and Their Interactions*

| Predictor | *df* | *χ2* | *p* |
|---|---|---|---|
| Rel | 2 | 125.88 | <.001*** |
| RT.cmc | 1 | 405.39 | <.001*** |
| Abs | 2 | 335.72 | <.001*** |
| Rel × Abs | 4 | 10.37 | .035* |

Note. Rel: Relative evidence; Abs: Absolute evidence;
RT.cmc: Response time (cluster-mean centered).
*p <.05 **p <.01 ***p <.001

Table S18

*Experiment 1 Regression Coefficients for Predicting Confidence (Error Trials) from Relative Evidence, RT, Absolute Evidence, and Their Interactions*

| Parameters | Estimate | *SE* | *z* | *p* |
|---|---|---|---|---|
| Intercept | 4.77 | 0.13 | 35.46 | <.001*** |
| Low Rel | 0.27 | 0.02 | 11.09 | <.001*** |
| Med Rel | -0.01 | 0.03 | -0.45 | .654 |
| RT.cmc | -0.00 | 0.00 | -20.39 | <.001*** |
| Low Abs | -0.52 | 0.03 | -17.07 | <.001*** |
| Med Abs | 0.12 | 0.03 | 4.72 | <.001*** |
| Low Rel × Low Abs | 0.04 | 0.04 | 1.09 | .275 |
| Med Rel × Low Abs | -0.03 | 0.04 | -0.85 | .393 |
| Low Rel × Med Abs | 0.06 | 0.03 | 1.75 | .079 |
| Med Rel × Med Abs | -0.01 | 0.04 | -0.24 | .807 |

Note. Intercept represents the estimate for high relative and high absolute evidence. Rel: Relative evidence; Abs: Absolute evidence; RT.cmc: Response time (cluster-mean centered).
*p <.05 **p <.01 ***p <.001

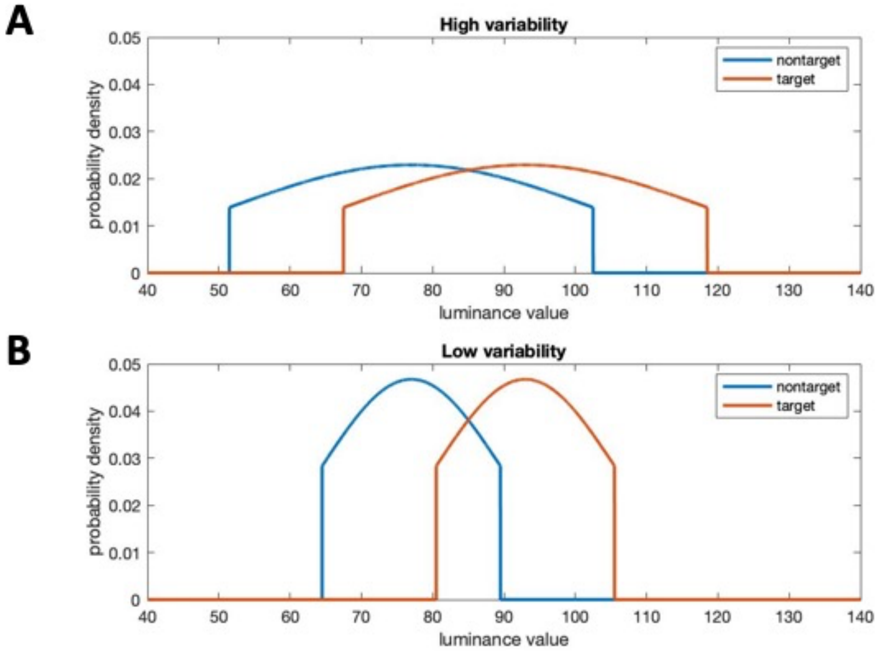

Figure S3. Example of luminance value distributions in (A) high variability conditions and (B) low variability conditions. Specifically, this example demonstrates the distributions when relative evidence is high and absolute evidence is low.

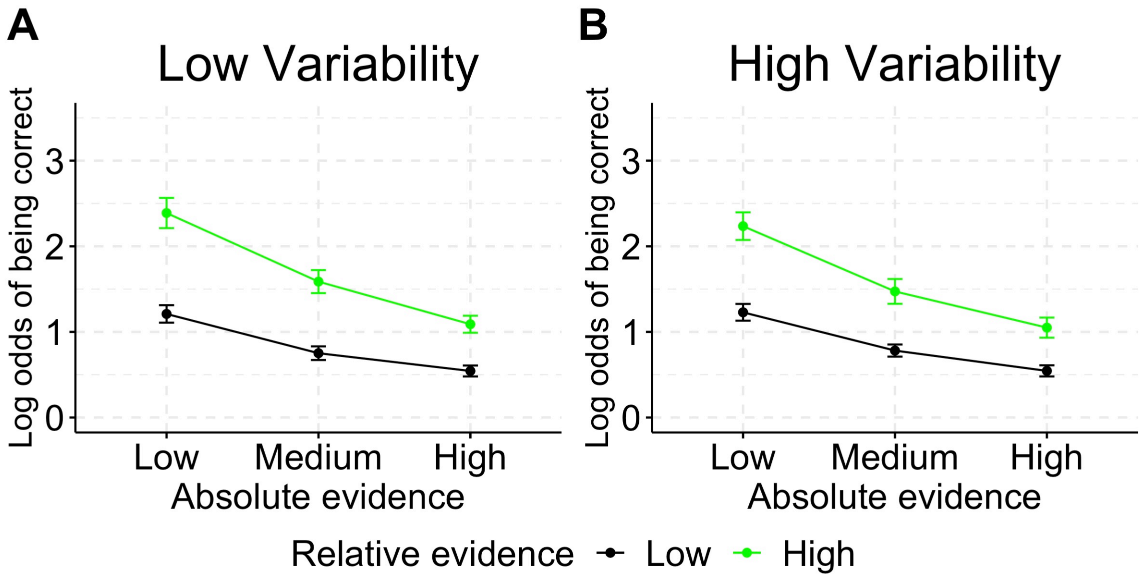

*Figure S4*. Experiment 2 mean log odds of being correct in each condition. Error bars represent SEM.

Table S19
*Mean luminance values for all experimental conditions of Experiment 2*

| | | Low absolute evidence | Medium absolute evidence | High absolute evidence |
|---|---|---|---|---|
| High luminance variability | High relative evidence | 77, 93 | 107, 123 | 137, 153 |
| | Low relative evidence | 77, 85 | 107, 115 | 137, 145 |
| Low luminance variability | High relative evidence | 77, 93 | 107, 123 | 137, 153 |
| | Low relative evidence | 77 85 | 107 115 | 137 145 |

Note. While mean luminance values were identical across high and low luminance variability conditions, high luminance conditions had standard deviations of 25.5, and low luminance conditions had standard deviations of 12.5.

**Accuracy (Log Odds of Being Correct)**

Table S20
*Experiment 2 Likelihood Ratio Tests Results for Predicting Accuracy (Log Odds of Being Correct) from Relative Evidence, Absolute Evidence, Luminance Variability, and Their Interactions*

| Predictor | *df* | $\chi 2$ | *p* |
|---|---|---|---|
| Rel | 1 | 576.22 | <.001*** |
| Abs | 2 | 588.29 | <.001*** |
| Var | 1 | 0.78 | .378 |
| Rel × Abs | 2 | 37.02 | <.001*** |
| Rel × Var | 1 | 2.77 | .096 |
| Abs × Var | 2 | 0.25 | .882 |
| Rel × Abs × Var | 2 | 0.53 | .767 |

Note. Rel: Relative evidence; Abs: Absolute evidence; Var: Luminance variability.
*p <.05 **p <.01 ***p <.001

Table S21
*Experiment 2 Regression Coefficients for Predicting Accuracy (Log Odds of Being Correct) from Relative Evidence, Absolute Evidence, Luminance Variability, and Their Interactions*

| Parameters | Estimate | *SE* | *z* | *p* |
|---|---|---|---|---|
| Intercept | 1.19 | 0.08 | 15.27 | <.001*** |
| Low Rel | -0.35 | 0.01 | -23.61 | <.001*** |
| Low Abs | 0.47 | 0.02 | 21.31 | <.001*** |
| Med Abs | -0.08 | 0.02 | -4.05 | <.001*** |
| Low Var | 0.01 | 0.01 | 0.88 | .378 |
| Low Rel × Low Abs | -0.12 | 0.02 | -5.35 | <.001*** |
| Low Rel × Med Abs | 0.02 | 0.02 | 0.75 | .456 |
| Low Rel × Low Var | -0.02 | 0.01 | -1.67 | .096 |
| Low Abs × Low Var | -0.01 | 0.02 | -0.41 | .680 |
| Med Abs × Low Var | 0.01 | 0.02 | 0.47 | .638 |
| Low Rel × Low Abs × Low Var | 0.01 | 0.02 | 0.29 | .770 |
| Low Rel × Med Abs × Low Var | -0.01 | 0.02 | -0.71 | .475 |

Note. Intercept represents the estimate for high relative and high absolute evidence. Rel: Relative evidence; Abs: Absolute evidence; Var: Luminance variability.
*p <.05 **p <.01 ***p <.001

**Response Time (Correct Trials)**

Table S22
*Experiment 2 Likelihood Ratio Tests Results for Predicting Response Time (Correct Trials) from Relative Evidence, Absolute Evidence, Luminance Variability, and Their Interactions*

| Predictor | *df* | $\chi 2$ | *p* |
|---|---|---|---|
| Rel | 1 | 106.53 | <.001*** |
| Abs | 2 | 44.46 | <.001*** |
| Var | 1 | 6.61 | .010* |
| Rel × Abs | 2 | 32.20 | <.001*** |
| Rel × Var | 1 | 0.37 | .541 |
| Abs × Var | 2 | 7.13 | .028* |
| Rel × Abs × Var | 2 | 6.24 | .044* |

Note. Rel: Relative evidence; Abs: Absolute evidence; Var: Luminance variability.
*p <.05 **p <.01 ***p <.001

Table S23

*Experiment 2 Regression Coefficients for Predicting Response Time (Correct Trials) from Relative Evidence, Absolute Evidence, Luminance Variability, and Their Interactions*

| Parameters | Estimate | *SE* | *z* | *p* |
|---|---|---|---|---|
| Intercept | 741.98 | 1.71 | 435.14 | <.001*** |
| Low Rel | 14.11 | 1.04 | 13.54 | <.001*** |
| Low Abs | 12.68 | 1.11 | 11.39 | <.001*** |
| Med Abs | -6.00 | 1.31 | -4.57 | <.001*** |
| Low Var | 3.52 | 1.09 | 3.22 | .001** |
| Low Rel × Low Abs | 10.44 | 1.33 | 7.85 | <.001*** |
| Low Rel × Med Abs | -7.55 | 1.42 | -5.33 | <.001*** |
| Low Rel × Low Var | 0.84 | 1.08 | 0.77 | .440 |
| Low Abs × Low Var | 0.06 | 1.21 | 0.05 | .964 |
| Med Abs × Low Var | 4.51 | 1.30 | 3.48 | .001** |
| Low Rel × Low Abs × Low Var | -1.89 | 1.15 | -1.64 | .100 |
| Low Rel × Med Abs × Low Var | 4.80 | 1.25 | 3.83 | <.001*** |

Note. Intercept represents the estimate for high relative and high absolute evidence. Rel: Relative evidence; Abs: Absolute evidence; Var: Luminance variability.
*p <.05 **p <.01 ***p <.001

**Response Time (Error Trials)**

Table S24

*Experiment 2 Likelihood Ratio Tests Results for Predicting Response Time (Error Trials) from Relative Evidence, Absolute Evidence, Luminance Variability, and Their Interactions*

| Predictor | *df* | $\chi 2$ | *p* |
|---|---|---|---|
| Rel | 1 | 23.74 | <.001*** |
| Abs | 2 | 37.22 | <.001*** |
| Var | 1 | 0.04 | .844 |
| Rel × Abs | 2 | 14.25 | <.001*** |
| Rel × Var | 1 | 0.16 | .685 |
| Abs × Var | 2 | 3.90 | .143 |
| Rel × Abs × Var | 2 | 3.37 | .185 |

Note. Rel: Relative evidence; Abs: Absolute evidence; Var: Luminance variability.
*p <.05 **p <.01 ***p <.001

Table S25

*Experiment 2 Regression Coefficients for Predicting Response Time (Error Trials) from Relative Evidence, Absolute Evidence, Luminance Variability, and Their Interactions*

| Parameters | Estimate | *SE* | *z* | *p* |
|---|---|---|---|---|
| Intercept | 784.84 | 4.05 | 193.66 | <.001*** |
| Low Rel | 12.86 | 2.20 | 5.86 | <.001*** |
| Low Abs | 23.00 | 2.82 | 8.16 | <.001*** |
| Med Abs | -6.64 | 2.33 | -2.85 | .004** |
| Low Var | 0.51 | 2.15 | 0.24 | .812 |
| Low Rel × Low Abs | 14.74 | 2.64 | 5.59 | <.001*** |
| Low Rel × Med Abs | -4.74 | 2.59 | -1.83 | .068 |
| Low Rel × Low Var | 1.06 | 2.27 | 0.47 | .641 |
| Low Abs × Low Var | 7.99 | 2.77 | 2.89 | .004** |
| Med Abs × Low Var | -4.44 | 2.68 | -1.65 | .098 |
| Low Rel × Low Abs × Low Var | -0.21 | 2.73 | -0.08 | .939 |
| Low Rel × Med Abs × Low Var | 5.29 | 2.79 | 1.89 | .058 |

Note. Intercept represents the estimate for high relative and high absolute evidence. Rel: Relative evidence; Abs: Absolute evidence; Var: Luminance variability.

*p <.05 **p <.01 ***p <.001

### Confidence (Correct Trials)

Table S26

*Experiment 2 Likelihood Ratio Tests Results for Predicting Confidence (Correct Trials) from Relative Evidence, Absolute Evidence, Luminance Variability, and Their Interactions*

| Predictor | *df* | $\chi 2$ | *p* |
|---|---|---|---|
| Rel | 1 | 261.57 | <.001*** |
| Abs | 2 | 191.90 | <.001*** |
| Var | 1 | 2.72 | .099 |
| Rel × Abs | 2 | 47.83 | <.001*** |
| Rel × Var | 1 | 0.45 | .501 |
| Abs × Var | 2 | 2.45 | .294 |
| Rel × Abs × Var | 2 | 1.15 | .561 |

Note. Rel: Relative evidence; Abs: Absolute evidence; Var: Luminance variability.
*p <.05 **p <.01 ***p <.001

Table S27

*Experiment 2 Regression Coefficients for Predicting Confidence (Correct Trials) from Relative Evidence, Absolute Evidence, Luminance Variability, and Their Interactions*

| Parameters | Estimate | *SE* | *z* | *p* |
|---|---|---|---|---|
| Intercept | 5.73 | 0.14 | 40.43 | <.001*** |
| Low Rel | -0.12 | 0.01 | -16.22 | <.001*** |
| Low Abs | -0.12 | 0.01 | -12.04 | <.001*** |
| Med Abs | -0.01 | 0.01 | -0.54 | .593 |
| Low Var | -0.01 | 0.01 | -1.65 | .099 |
| Low Rel × Low Abs | -0.07 | 0.01 | -6.79 | <.001*** |
| Low Rel × Med Abs | 0.02 | 0.01 | 1.97 | .048* |
| Low Rel × Low Var | -0.00 | 0.01 | -0.67 | .501 |
| Low Abs × Low Var | -0.01 | 0.01 | -1.00 | .316 |
| Med Abs × Low Var | -0.01 | 0.01 | -0.60 | .550 |
| Low Rel × Low Abs × Low Var | 0.00 | 0.01 | 0.11 | .909 |
| Low Rel × Med Abs × Low Var | -0.01 | 0.01 | -1.00 | .318 |

Note. Intercept represents the estimate for high relative and high absolute evidence. Rel: Relative evidence; Abs: Absolute evidence; Var: Luminance variability.

*p <.05 **p <.01 ***p <.001

**Confidence (Error Trials)**

Table S28

*Experiment 2 Likelihood Ratio Tests Results for Predicting Confidence (Error Trials) from Relative Evidence, Absolute Evidence, Luminance Variability, and Their Interactions*

| Predictor | *df* | $\chi2$ | *p* |
|---|---|---|---|
| Rel | 1 | 57.71 | <.001*** |
| Abs | 2 | 400.02 | <.001*** |
| Var | 1 | 0.85 | .355 |
| Rel × Abs | 2 | 2.65 | .265 |
| Rel × Var | 1 | 4.35 | .037* |
| Abs × Var | 2 | 8.47 | .014* |
| Rel × Abs × Var | 2 | 3.79 | .150 |

Note. Rel: Relative evidence; Abs: Absolute evidence; Var: Luminance variability.
*p <.05 **p <.01 ***p <.001

Table S29

*Experiment 2 Regression Coefficients for Predicting Confidence (Error Trials) from Relative Evidence, Absolute Evidence, Luminance Variability, and Their Interactions*

| Parameters | Estimate | *SE* | *z* | *p* |
|---|---|---|---|---|
| Intercept | 4.78 | 0.13 | 35.64 | <.001*** |
| Low Rel | 0.14 | 0.02 | 7.61 | <.001*** |
| Low Abs | -0.52 | 0.03 | -17.88 | <.001*** |
| Med Abs | 0.08 | 0.03 | 3.27 | .001** |
| Low Var | -0.02 | 0.02 | -0.92 | .355 |
| Low Rel × Low Abs | 0.03 | 0.03 | 1.16 | .245 |
| Low Rel × Med Abs | 0.00 | 0.03 | 0.20 | .845 |
| Low Rel × Low Var | 0.04 | 0.02 | 2.09 | .037* |
| Low Abs × Low Var | -0.08 | 0.03 | -2.73 | .006** |
| Med Abs × Low Var | 0.06 | 0.03 | 2.46 | .014* |
| Low Rel × Low Abs × Low Var | 0.05 | 0.03 | 1.90 | .057 |
| Low Rel × Med Abs × Low Var | -0.02 | 0.03 | -0.83 | .408 |

Note. Intercept represents the estimate for high relative and high absolute evidence. Rel: Relative evidence; Abs: Absolute evidence; Var: Luminance variability.

*p <.05 **p <.01 ***p <.001

**Change of Mind (Log Odds of Confidence Lower Than 4) in Correct Trials**

Table S30

*Experiment 2 Likelihood Ratio Tests Results for Predicting Change of Mind (Log Odds of Confidence Lower Than 4) in Correct Trials from Relative Evidence, Absolute Evidence, Luminance Variability, and Their Interactions*

| Predictor | *df* | $\chi 2$ | *p* |
|---|---|---|---|
| Rel | 1 | 36.33 | <.001*** |
| Abs | 2 | 5.26 | .072 |
| Var | 1 | 0.27 | .601 |
| Rel × Abs | 2 | 9.72 | .008** |
| Rel × Var | 1 | 0.08 | .780 |
| Abs × Var | 2 | 0.49 | .784 |
| Rel × Abs × Var | 2 | 0.33 | .846 |

Note. Rel: Relative evidence; Abs: Absolute evidence; Var: Luminance variability.
*p <.05 **p <.01 ***p <.001

Table S31

*Experiment 2 Regression Coefficients for Predicting Change of Mind (Log Odds of Confidence Lower Than 4) in Correct Trials from Relative Evidence, Absolute Evidence, Luminance Variability, and Their Interactions*

| Parameters | Estimate | *SE* | *z* | *p* |
|---|---|---|---|---|
| Intercept | 3.55 | 0.27 | 13.30 | <.001*** |
| Low Rel | -0.19 | 0.03 | -6.05 | <.001*** |
| Low Abs | -0.10 | 0.04 | -2.29 | .022* |
| Med Abs | 0.03 | 0.04 | 0.77 | .439 |
| Low Var | 0.02 | 0.03 | 0.53 | .599 |
| Low Rel × Low Abs | -0.13 | 0.04 | -3.12 | .002** |
| Low Rel × Med Abs | 0.05 | 0.04 | 1.22 | .224 |
| Low Rel × Low Var | 0.01 | 0.03 | 0.28 | .779 |
| Low Abs × Low Var | 0.03 | 0.04 | 0.66 | .510 |
| Med Abs × Low Var | -0.02 | 0.04 | -0.51 | .607 |
| Low Rel × Low Abs × Low Var | -0.02 | 0.04 | -0.53 | .595 |
| Low Rel × Med Abs × Low Var | 0.00 | 0.04 | 0.04 | .971 |

Note. Intercept represents the estimate for high relative and high absolute evidence. Rel: Relative evidence; Abs: Absolute evidence; Var: Luminance variability.

*p <.05 **p <.01 ***p <.001

**Change of Mind (Log Odds of Confidence Lower Than 4) in Error Trials**

Table S32

*Experiment 2 Likelihood Ratio Tests Results for Predicting Change of Mind (Log Odds of Confidence Lower Than 4) in Error Trials from Relative Evidence, Absolute Evidence, Luminance Variability, and Their Interactions*

| Predictor | *df* | *χ2* | *p* |
|---|---|---|---|
| Rel | 1 | 51.07 | <.001*** |
| Abs | 2 | 179.30 | <.001*** |
| Var | 1 | 0.00 | .976 |
| Rel × Abs | 2 | 3.62 | .164 |
| Rel × Var | 1 | 3.66 | .056 |
| Abs × Var | 2 | 7.92 | .019* |
| Rel × Abs × Var | 2 | 7.20 | .027* |

Note. Rel: Relative evidence; Abs: Absolute evidence; Var: Luminance variability.
*p <.05 **p <.01 ***p <.001

Table S33
*Experiment 2 Regression Coefficients for Predicting Change of Mind (Log Odds of Confidence Lower Than 4) in Error Trials from Relative Evidence, Absolute Evidence, Luminance Variability, and Their Interactions*

| Parameters | Estimate | *SE* | *z* | *p* |
|---|---|---|---|---|
| Intercept | 1.57 | 0.18 | 8.81 | <.001*** |
| Low Rel | 0.24 | 0.03 | 7.18 | <.001*** |
| Low Abs | -0.61 | 0.05 | -12.22 | <.001*** |
| Med Abs | 0.08 | 0.05 | 1.75 | .081 |
| Low Var | 0.00 | 0.03 | 0.03 | .976 |
| Low Rel × Low Abs | 0.07 | 0.05 | 1.45 | .147 |
| Low Rel × Med Abs | 0.01 | 0.05 | 0.30 | .767 |
| Low Rel × Low Var | 0.06 | 0.03 | 1.92 | .055 |
| Low Abs × Low Var | -0.14 | 0.05 | -2.81 | .005** |
| Med Abs × Low Var | 0.07 | 0.05 | 1.58 | .113 |
| Low Rel × Low Abs × Low Var | 0.12 | 0.05 | 2.53 | .011* |
| Low Rel × Med Abs × Low Var | -0.03 | 0.05 | -0.55 | .582 |

Note. Intercept represents the estimate for high relative and high absolute evidence. Rel: Relative evidence; Abs: Absolute evidence; Var: Luminance variability.
*p <.05 **p <.01 ***p <.001

**Confidence (Correct Trials)**

Table S34

*Experiment 2 Likelihood Ratio Tests Results for Predicting Confidence (Correct Trials) from Relative Evidence, Luminance Variability, RT, Absolute Evidence, and Their Interactions*

| Predictor | *df* | $\chi 2$ | *p* |
|---|---|---|---|
| Rel | 1 | 195.23 | <.001*** |
| Var | 1 | 1.08 | .299 |
| RT.cmc | 1 | 1044.45 | <.001*** |
| Abs | 2 | 169.31 | <.001*** |
| Rel × Var | 1 | 0.24 | .622 |
| Rel × Abs | 2 | 33.46 | <.001*** |
| Var × Abs | 2 | 1.81 | .405 |
| Rel × Var × Abs | 2 | 0.54 | .765 |

Note. Rel: Relative evidence; Abs: Absolute evidence; Var: Luminance variability; RT.cmc: Response time (cluster-mean centered).
*p <.05 **p <.01 ***p <.001

Table S35

*Experiment 2 Regression Coefficients for Predicting Confidence (Correct Trials) from Relative Evidence, Luminance Variability, RT, Absolute Evidence, and Their Interactions*

| Parameters | Estimate | *SE* | *z* | *p* |
|---|---|---|---|---|
| Intercept | 5.73 | 0.14 | 40.43 | <.001*** |
| Low Rel | -0.10 | 0.01 | -14.00 | <.001*** |
| Low Var | -0.01 | 0.01 | -1.04 | .299 |
| RT.cmc | -0.00 | 0.00 | -32.72 | <.001*** |
| Low Abs | -0.11 | 0.01 | -10.95 | <.001*** |
| Med Abs | -0.01 | 0.01 | -1.18 | .238 |
| Low Rel × Low Var | -0.00 | 0.01 | -0.49 | .622 |
| Low Rel × Low Abs | -0.05 | 0.01 | -5.55 | <.001*** |
| Low Rel × Med Abs | 0.01 | 0.01 | 1.13 | .258 |
| Low Var × Low Abs | -0.01 | 0.01 | -1.06 | .287 |
| Low Var × Med Abs | -0.00 | 0.01 | -0.23 | .815 |
| Low Rel × Low Var × Low Abs | -0.00 | 0.01 | -0.07 | .942 |
| Low Rel × Low Var × Med Abs | -0.01 | 0.01 | -0.61 | .541 |

Note. Intercept represents the estimate for high relative evidence, high absolute evidence, and high luminance variability. Rel: Relative evidence; Abs: Absolute evidence; Var: Luminance variability; RT.cmc: Response time (cluster-mean centered).

*p <.05 **p <.01 ***p <.001

**Confidence (Error Trials)**

Table S36

*Experiment 2 Likelihood Ratio Tests Results for Predicting Confidence (Error Trials) from Relative Evidence, Luminance Variability, RT, Absolute Evidence, and Their Interactions*

| Predictor | *df* | $\chi 2$ | *p* |
|---|---|---|---|
| Rel | 1 | 75.31 | <.001*** |
| Var | 1 | 0.66 | .418 |
| RT.cmc | 1 | 357.34 | <.001*** |
| Abs | 2 | 357.69 | <.001*** |
| Rel × Var | 1 | 4.95 | .026* |
| Rel × Abs | 2 | 5.87 | .053 |
| Var × Abs | 2 | 6.83 | .033* |
| Rel × Var × Abs | 2 | 4.66 | .097 |

Note. Rel: Relative evidence; Abs: Absolute evidence; Var: Luminance variability; RT.cmc: Response time (cluster-mean centered).
*p <.05 **p <.01 ***p <.001

Table S37

*Experiment 2 Regression Coefficients for Predicting Confidence (Error Trials) from Relative Evidence, Luminance Variability, RT, Absolute Evidence, and Their Interactions*

| Parameters | Estimate | *SE* | *z* | *p* |
|---|---|---|---|---|
| Intercept | 4.78 | 0.13 | 35.64 | <.001*** |
| Low Rel | 0.16 | 0.02 | 8.70 | <.001*** |
| Low Var | -0.01 | 0.02 | -0.81 | .418 |
| RT.cmc | -0.00 | 0.00 | -19.14 | <.001*** |
| Low Abs | -0.47 | 0.03 | -16.80 | <.001*** |
| Med Abs | 0.07 | 0.02 | 2.90 | .004** |
| Low Rel × Low Var | 0.04 | 0.02 | 2.23 | .026* |
| Low Rel × Low Abs | 0.05 | 0.03 | 1.88 | .060 |
| Low Rel × Med Abs | 0.00 | 0.02 | 0.06 | .951 |
| Low Var × Low Abs | -0.07 | 0.03 | -2.43 | .015* |
| Low Var × Med Abs | 0.06 | 0.02 | 2.25 | .024* |
| Low Rel × Low Var × Low Abs | 0.06 | 0.03 | 2.04 | .042* |
| Low Rel × Low Var × Med Abs | -0.02 | 0.02 | -0.67 | .502 |

Note. Intercept represents the estimate for high relative evidence, high absolute evidence, and high luminance variability. Rel: Relative evidence; Abs: Absolute evidence; Var: Luminance variability; RT.cmc: Response time (cluster-mean centered).

*p <.05 **p <.01 ***p <.001
